# Tracking Single Particles for Hours via Continuous DNA-mediated Fluorophore Exchange

**DOI:** 10.1101/2020.05.17.100354

**Authors:** Johannes Stein, Florian Stehr, Julian Bauer, Christian Niederauer, Ralf Jungmann, Kristina Ganzinger, Petra Schwille

## Abstract

Fluorophores are commonly used to covalently label biomolecules for monitoring their motion in single particle tracking experiments. However, photobleaching is still a major bottleneck in these experiments, as the fluorophores’ finite photon budget typically limits observation times to merely a few seconds. Here, we overcome this inherent constraint *via* continuous fluorophore exchange based on DNA-PAINT, whereby fluorescently-labeled oligonucleotides bind to a 54 bp single-stranded DNA handle attached to the molecule of interest. When we assayed our approach *in vitro* by tracking single DNA origami, first surface-immobilized and subsequently diffusing on supported lipid bilayers, we were able to observe these origami for up to hours without losing their fluorescence signals. Our versatile and easily implemented labeling approach allows monitoring single-molecule motion and interactions over unprecedented observation periods, opening the doors to advanced quantitative studies.

## Introduction

If one could make three wishes for a fluorescent label for Single Particle Tracking (SPT), one would ask for 1) an infinitely small and non-invasive label that is 2) so bright that we could follow it by plain eyesight for an infinite amount of time and that 3) could be specifically attached (1:1) to the target particle of interest. Obviously, these demands are not only unrealistic but also, in part, mutually exclusive. Established labeling strategies, therefore, make trade-offs, and are thus either better performing in one of those aspects or another. Organic dyes, for instance, are the smallest possible choice and offer specific 1:1 labeling, but as their photon budget is limited, particles can typically not be observed for longer periods than tens of seconds. Chemical compounds, such as oxygen scavenging systems and triplet state quenchers, can improve the fluorescence performance of organic dyes^1–5^, but they are not compatible with live cell experiments as oxygen is required for respiratory-chain reactions^6^. On the other end of the size spectrum, Quantum Dots (QDs) have become a popular fluorescent label for SPT experiments due to their superior fluorescence properties with respect to brightness and photobleaching compared to organic dyes or fluorescent proteins^7^. However, biocompatible QDs are up to orders of magnitude larger than dyes/proteins, potentially impairing the dynamics of the biological system of interest^7,8^, subject to photoblinking^8^ and furthermore difficult to functionalize at the desired 1:1 stoichiometry^8^, resulting in the (possibly unknown) labeling of multiple molecules and thus experimental artefacts. More recently, researchers have developed novel label designs^9^ or labeling strategies^10^ reporting drastic improvements on some of the above-stated demands.

Here, we introduce a live cell compatible labeling approach for SPT based on DNA-mediated fluorophore exchange which not only provides superior fluorescence properties compared to individual organic dyes, but is smaller in size than QDs and allows 1:1 labeling. This labeling strategy allowed us to continuously observe fixed single particles for up to one hour, and mobile particles for up to 18 minutes, thus for durations that are several orders of magnitude longer than those achieved for single organic dyes and even exceed those routinely achieved with QDs, enabling us to sample diffusion properties across the entire field-of-view.

## Results

### DNA-mediated fluorophore exchange for creating a long-lived fluorescent label

The principle of our labeling approach relies on that of DNA-PAINT^11^ (Points Accumulation for Imaging in Nanoscale Topography): the continuous binding, detachment and re-binding of single-stranded and fluorescently-labeled oligonucleotides (‘imagers’) to a complementary single DNA strand on the molecule of interest, which we will refer to as ‘tracking handle’ (TH) (**Figure 1**a). The 54 base pairs (bp) TH sequence is composed of 18× repetitions of the codon ‘CTC’ and both the 3’ and the 5’ TH end can be modified with functional groups for the desired chemical attachment strategy to the target molecule (e.g. click-chemistry, SNAP-tag, HALO-tag etc.). Following DNA-PAINT, individual freely-diffusing imagers (8 bp; sequence: 5’-GAGGAGGA-3’-Cy3B) bind to the TH *via* reversible DNA hybridization of their short complementary oligonucleotide part (**Figure 1**b). Importantly, we designed the TH-imager sequence in such a way that at least one emitting imager is bound to the TH at all times, while exchange is mediated by a continuous turnover of ‘fresh’ imagers from solution. In other words, before the last bound imager dissociates, or before its dye molecule photobleaches, ideally a new imager has already arrived and hybridized to the TH, theoretically producing an infinitely long fluorescence signal from the TH. To achieve this, we allowed multiple imagers to bind simultaneously (max. 6 imagers at once) to the TH (green marked regions in sequence in **Figure 1**a). Second, we strived to maximize the imager association rate (*k*_on_) in order to ensure a fast turnover^12,13^. Third, we adjusted the dwell times (*k*_off_) of imagers bound to the TH with respect to the photon budget of Cy3B dyes such that imagers unbind before they photobleach (to prevent bleached imagers blocking TH binding sites). **Figure 1**c gives an overview of the labeling principle, schematically illustrating an idealized fluorescence signal originating from a TH attached as label to a target molecule. First, three imagers are simultaneously bound to the TH, before one of the imagers dissociates resulting in a drop in the fluorescence signal (i). Next, the dye molecule of one of the remaining two imagers photobleaches (ii), equally resulting in an intensity drop. Before the last imager dissociates or photobleaches (which would imply a loss of the fluorescence signal, i.e. the position of the target particle), a fresh imager strand binds to the TH (iii) resulting in an intensity increase. However, while theoretically infinite observation times of the TH are possible, this would come at the cost of an increased length of the TH, potentially altering the diffusion properties of the particle of interest compared to single-dye labeling, or at the cost of increased imager concentrations, causing a high fluorescence background. At 54bp, our tracking handle is too short to allow for unlimited observations times, but is still smaller in size than most conventionally used QDs (see **Supplementary Figure 1** for a size comparison true to scale).

**Figure 1.**
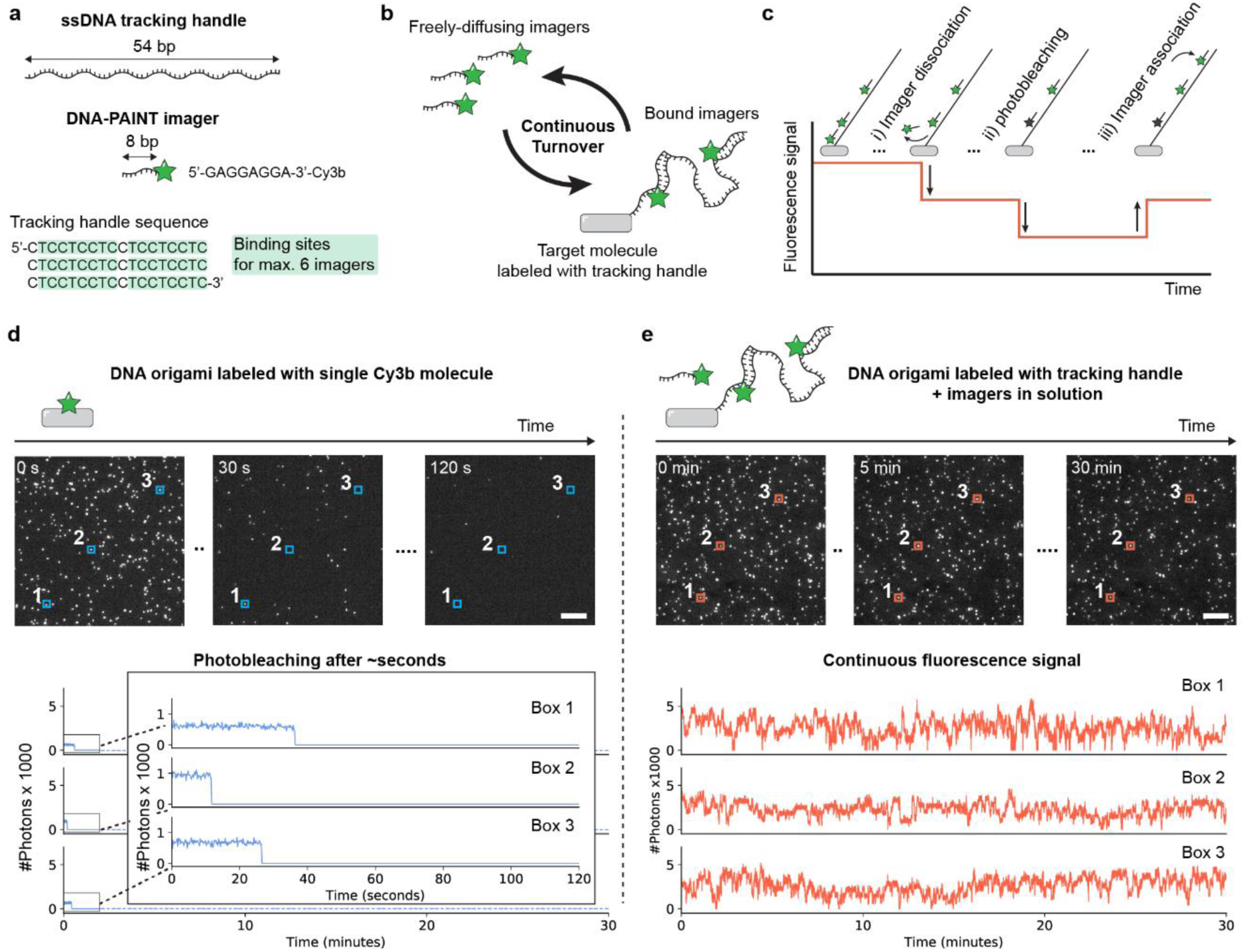
DNA-mediated continuous fluorophore exchange. (**a**) Design of the 54 base-pair ssDNA tracking handle (TH) and DNA-PAINT imagers. (**b**) Principle of repurposing DNA-PAINT for single-particle tracking experiments. Freely-diffusion imagers are binding and unbinding in a continuous turnover to the TH attached to the target molecule. (**c**) Schematic of intensity fluctuations recorded from a TH caused by imager dissociation, association and photobleaching. (**d**) TIRFM imaging of static single-dye labeled DNA origami. Localized fluorescence traces end abruptly when photobleaching occurs. (e) TIRFM imaging of static DNA origami labeled with the TH and freely-diffusing imagers in the solution. The fluorescence signal exhibits continuous intensity fluctuations due to imager exchange. Scale bars, 5 *μ*m in (d,e).

To demonstrate the improved fluorescence properties of the TH compared to a single dye molecule as a label for SPT, we compared the photobleaching behavior of single Cy3B molecules attached to static DNA origami to that of origami labeled with the TH as visualized by Cy3B imagers. To do so, we recorded 2-minute image sequences of surface-immobilized and well-separated DNA origami labeled by single Cy3B molecules (hereafter referred to as ‘single-dye origami’ or SD origami) via TIRFM^14^ (Total Internal Reflection Fluorescence Microscopy) to monitor the photobleaching behavior of single Cy3B molecules over time (**Figure 1**d, top, see **Supplementary Table 1** for detailed imaging parameters of all presented data). The localized fluorescence vs. time traces (referred to as ‘fluorescence trace’ in the following) of three exemplary dyes corresponding to the blue three boxes in the top images illustrate that bleaching occurs on the time scale of tens of seconds (**Figure 1**d, bottom) and after 2 min nearly all dyes had entered a permanently dark state. In contrast, when we recorded a 30-minute image sequence of a sample containing the same surface-immobilized DNA origami, but this time labeled with THs (‘TH origami’) and additionally 40 nM of imagers added to the solution, a large fraction of all TH origami was still observable *even after 30 minutes* under identical acquisition conditions. (**Figure 1**e, top). In line with this visual impression, the three exemplary TH fluorescence traces (orange) reveal fluctuating but continuous intensity signals over the course of 30 minutes (**Figure 1**e, bottom). For an objective and quantitative comparison of the fluorescence properties between single dyes and THs, we identified three key measures relevant for SPT: i) the duration of each particle trajectory (for a more precise MSD description^15^), ii) the total number of collectable trajectories per particle (for larger statistics) and iii) the fluorescence brightness of the label (for a higher localization precision^16^). In the following section, we introduce the reader to our key analysis metrics based on those three quality measures, using the examples of the two data sets shown in **Figure 1**d & e.

### Key metrics for an objective and quantitative comparison of the fluorescence properties between single dyes and THs

Both immobilized DNA origami imaging data sets (both for SD origami and TH origami) were subjected to the same post-processing pipeline. First, a standard single molecule localization algorithm^11^ was applied to obtain a pointillist super-resolution image of the DNA origami appearing as localization clusters. Subsequently, we automatically detected all localization clusters as previously described^17^ and extracted the corresponding fluorescence traces (see **Supplementary Figure 2**). A typical fluorescence trace as recorded from a SD origami consisted of a single fluorescence burst at the start of image acquisition with an abrupt ending caused by photobleaching, as depicted in **Figure 2**a (example trace of SD941 arbitrarily selected out of all ∼3,300 SD origami in the data set surpassing the filter criteria described in **Supplementary Figure 3**). Neglecting short interruptions, e.g. due to photoblinking or signal-to-noise variations, the trajectory duration *τ*_1_ of a single dye was determined by its photostability. Hence, in the language of SPT, regardless of the measurement duration, we were only able to record one trajectory per SD origami. Note that for all surface-immobilized experiments (for both SD origami and TH origami) we ignored interruptions in the fluorescence trace of just a single frame when determining the trajectory durations (see ‘ignore’ in **Supplementary Figure 2**).

**Figure 2.**
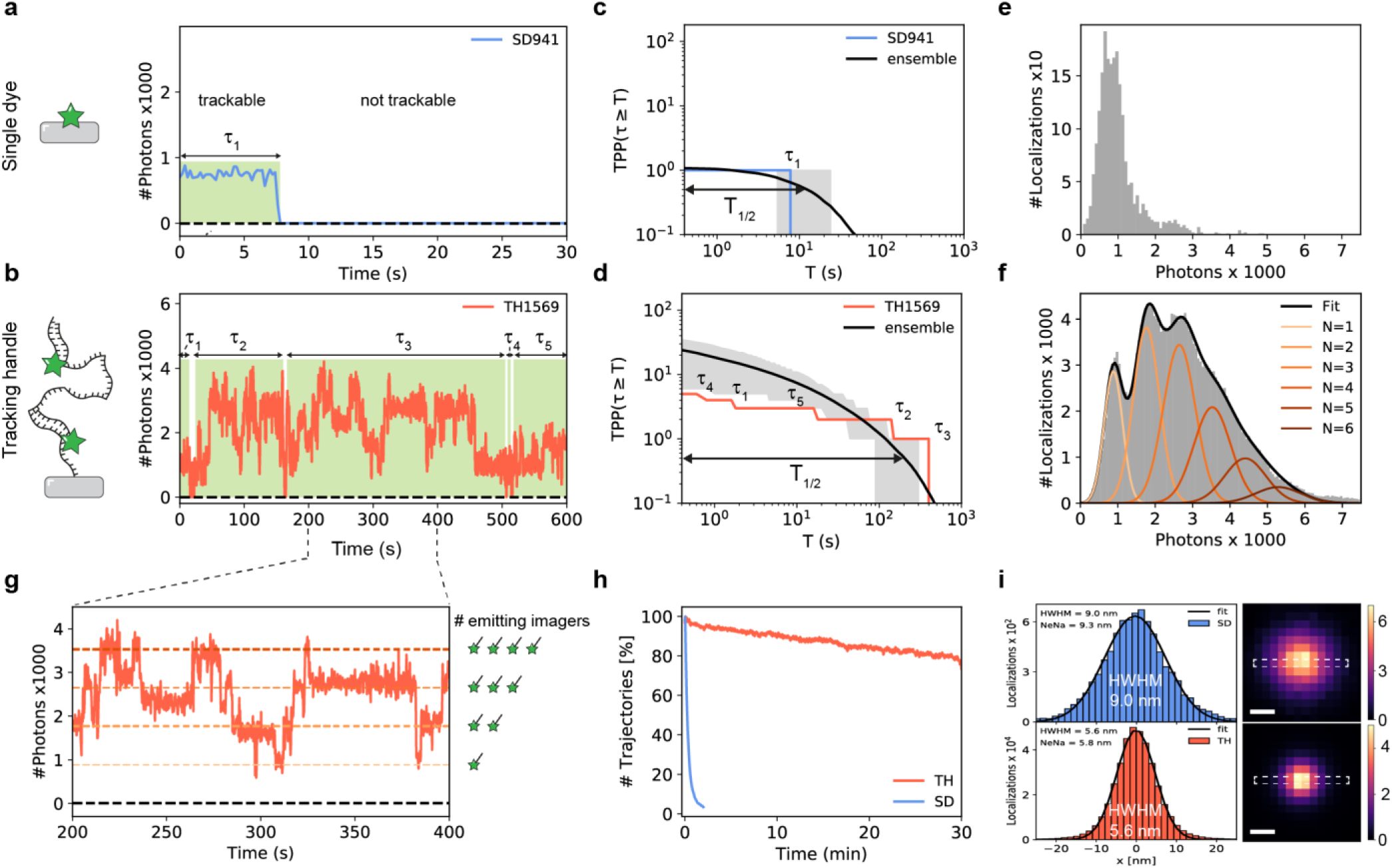
Comparing the fluorescence properties of SD origami vs TH origami. (**a**) SD example fluorescence trace. Typically, a SD origami produces one fluorescence burst resulting in a single trajectory. (**b**) TH example fluorescence trace. Typically, a TH origami exhibits fluctuating fluorescence intensity with short interruptions. (**c**) Plot of *TPP*(*τ* ≥ *T*) vs. *T* for the SD fluorescence trace in (a) (blue) and averaged over all SD origami in data set (black). (**d**) Plot of *TPP*(*τ* ≥ *T*) vs. *T* for the TH fluorescence trace in (b) (orange) and averaged over all TH origami in data set (black). (**e**) Number of photons detected per localization for SD origami showing a unimodal distribution. (**f**) Number of photons detected per localization for TH origami showing a multimodal distribution. A fit (black) consisting of the sum of 6 Gaussians (orange) was applied to analyze how many emitting imagers were present over time. (**g**) Zoom-in of TH fluorescence trace from (b) revealing step like behavior and the number of currently emitting imagers bound to the TH. The dashed horizontal lines indicate the center of the 6 Gaussians obtained in (f). (**h**) Number of trajectories per frame (i.e. emitting labels) vs. measurement time normalized to initial trajectory number. (**i**) Increased localization precision for TH. Cross-sectional histograms over indicated regions (white dashed box) in averaged super-resolved images of immobilized SD origami (top, blue) and TH origami (bottom, orange). Scale bars, 10 nm in (i). Error in (c,d) refers to interquartile range, indicated as grey shaded area.

A typical fluorescence trace of a TH origami, in contrast, showed an almost continuous signal with fluctuating intensity as the number of bound imagers varied over time (**Figure 2**b; TH1569 selected out of all ∼2,600 clusters in the data set after filtering; a 10-minute subset of the trace is displayed for illustration purposes). However, we also observed short interruptions in TH fluorescence traces of a few frames and in the depicted example, this resulted in total in five trajectories of durations *τ*_1−5_ in the range of ∼10-200 s. These interruptions originated from the stochastic nature of imager binding and photobleaching causing the TH to enter a short non-fluorescent state. Comparing both fluorescence traces in **Figure 2**a and b, one can deduce the improvement of the TH with respect to the three previously defined quality measures by eye: the TH handle produces i) more trajectories, which on average have ii) a longer duration and iii) a higher fluorescence brightness. Note that in contrast to permanent labeling with a single dye for the TH the number of recorded trajectories per particle (*TPP*) will grow with the duration of the measurement.

Given a certain number of particles within a field of view (FOV), the SPT-experimentalist aims to maximize the number of long trajectories within the measurement time^8^. From the fluorescence trace of every immobilized DNA origami *i* in a data set (*i* = 1,2, …, *M*, where *M* denotes the total number of origami after filtering) we therefore calculated the number of trajectories per particle *TPP*_*i*_ (*τ* ≥ *T*) with a duration *τ* longer or equal to time *T* (**Figure 2**c and d). The blue curve in **Figure 2**c displays the resulting *TPP*_*SD94*1_(*τ* ≥ *T*) vs. *T* plot based on fluorescence trace SD941 shown in **Figure 2**a. As expected for a single dye, a single trajectory is produced and thus *TPP*_*SD94*1_(*τ* ≥ *T*) is equal to one, as long as *T* does not exceed the trajectory duration *τ*_1_ and drops to zero for higher values of *T*. Calculation of *TPP*_*i*_ (*τ* ≥ *T*) for all *M* fluorescence traces of the data set and subsequent averaging yielded the ensemble mean 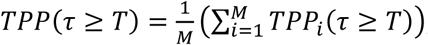, illustrated as black line with its interquartile range as grey contour (**Figure 2**c). We defined the characteristic half-life time *T*_1/2_ as the time at which the ensemble mean falls below one-half (i.e. *TPP*(*τ* ≥ *T*_1/2_) = 0.5). Hence, *TPP*(*τ* ≥ *T*) simultaneously allows a quantitative description of how many trajectories one can expect per particle/origami and the expected average trajectory duration. This becomes explicitly clear when interpreting the ensemble curve *TPP*(*τ* ≥ *T*) in **Figure 2**c in simpler words: All ∼3,000 SD origami in the data set produced on average a single trajectory (see *y*-axis intercept at 1) and half of these (i.e. ∼1,500) had a duration of at least 11 s.

We can now apply the same reasoning for the TH origami data set (note that for illustration purposes we applied the analysis on a 10-minute subset of the data). The orange curve in **Figure 2**d shows *TPP*_*TH*1569_(*τ* ≥ *T*) corresponding to the five trajectories *τ*_1,…,5_ of the individual fluorescence trace TH1569 (see **Figure 2**b) showing the step-like integer decrease at the end of trajectory duration (i.e. the curve starts at 5 for short *T* and finally drops to 0 at the end of the longest trajectory *τ*_3_). Again, the black curve corresponds to the ensemble mean *TPP*(*τ* ≥ *T*) and the grey contour to the interquartile range (number TH origami after filtering: *M* ∼2,500). Each TH yielded on average ∼22 trajectories over the measurement duration of 10 minutes and *T*_1/2_ analysis revealed that we registered 1,250 trajectories with a duration of at least 200 s (>3 minutes), resulting in an increase in both the number of tracks and in *T*_1/2_ of a factor of ∼20x compared to SD origami.

The TH is designed to simultaneously host a maximum of 6 imagers, but the actual number of imagers bound at any point in time is governed by the reaction rates *k*_on_ and *k*_off_ and will therefore vary due to the stochastic nature of those reactions. To assess the number of simultaneously bound imagers we extracted the detected number of photons per localization. We assumed a linear increase in fluorescence intensity with every fresh imager binding and analogously a linear decrease with every imager unbinding or photobleaching of the dye. First, we analyzed the photon counts for the SD origami data set, which exhibited a single peak indicating that under the applied acquisition conditions we detected on average around 900 photons per localization from a single Cy3B molecule (see **Figure 2**e). In order to account for imaging artifacts causing variations in the number of detected photons^18^, we only used origami lying within the central circular region of the FOV (diameter = 200 px). Subsequently, we performed the same photon count analysis on the TH data set, which resulted in a multimodal distribution with distinct peaks that seemed to be in an equidistant spacing and to be located at multiples of the lowest peak’s center value that coincides with that of the single peak from the SD origami data set. (**Figure 2**f compare to **Figure 2**e). This suggested that the observation of higher order photon counts could indeed be used to identify the number of emitting imagers (carrying single dyes) bound to a TH. To quantify the probability of how many imager strands are bound to the TH we fitted its photon count distribution with the sum (black) of 6 Gaussian functions *g*_i_ (colored) of the form 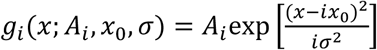 with freely floating amplitudes *A*_*i*_ for each *g*_*i*_ but global parameters *x*_0_ and *σ* for the sum, corresponding to the center and width of the lowest order peak (see **Figure 2**f, orange lines). The amplitudes *A*_*i*_ can directly be translated into a probability of *i* emitting imager strands being bound to the TH. Plotting the obtained photon levels *ix*_0_ on top of the fluorescence trace of **Figure 2**b (zoom) shows that indeed the fluorescence fluctuations follow a step-like behavior dependent on the number of emitting imagers bound at every time point (**Figure 2**g, for photon count histogram see **Supplementary Figure 4**).

Next, we compared the number of trajectories per frame normalized to the first frame and found that over the course of 30 min in each frame more than 80 % of all THs were detected (**Figure 2**h, orange curve). While ∼97 % of the SD origami photobleached after 120 s (**Figure 2**h, blue curve), also TH origami showed a 20 % decrease over 30 min due to photo-induced damage to the DNA caused by reactive oxygen species^19^.

Finally, we analyzed the effect of the previously described increased brightness of the TH compared to SD origami with respect to the localization precision. **Figure 2**i depicts cross-sectional histograms through the aligned and averaged images of several hundreds of both SD origami (top image and blue histogram) and TH origami (bottom image and orange histogram). The visual impression of sharper localization distribution for THs compared to SDs is confirmed by comparing the half width half maximum (HWHM) of the Gaussian fits to the two histograms (5.6 nm and 9.0 nm, respectively), which are in good agreement with the localization precision results based on Nearest-Neighbor Analysis^20^ (NeNA; 5.8 nm and 9.3 nm, respectively).

### What keeps the Tracking Handle (from) running

As a next step, we investigated the various factors determining the function and performance of the TH, again under ideal surface-immobilized conditions. The four main kinetic rates that control continuous imager exchange are: i) the rate of photobleaching *k*_photobleaching_, ii) the rate of photo-induced damage *k*_photodamage_, iii) the effective imager association rate *k*_on_ and iv) the dissociation rate *k*_off_ (**Figure 3**a). Depending on the experimental conditions, such as the temperature or the imaging buffer composition as well as the excitation laser power (irradiance) at which imaging is performed, either of these rates can play a more or less dominant role for the TH. We optimized the experimental conditions for SPT experiments in a live-cell compatible buffer L (see methods), at temperature T=21 °C and at an imager concentration of 40 nM (if data was acquired at deviating conditions this is explicitly stated).

**Figure 3.**
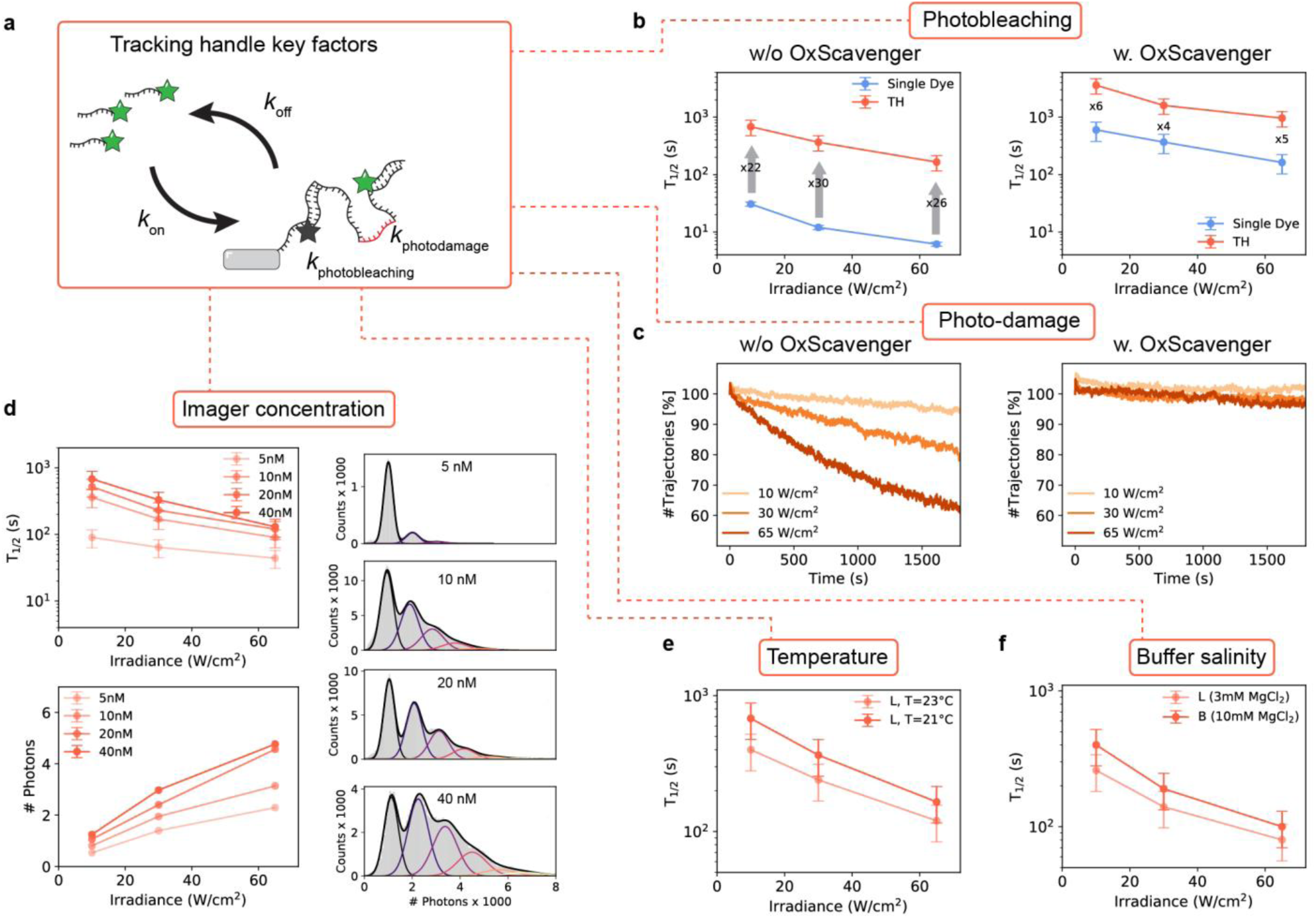
Tracking handle key factors. (**a**) Main rates determining the TH function (**b**) *T*_1/2_ vs. irradiance plots for SD origami (blue) and TH origami (orange) imaged at varying irradiances. Left and right panels show the case without and with using an oxygen scavenging system, respectively. Arrows indicates factors of increase of TH vs SD. (**c**) Number of trajectories per frame (i.e. emitting labels) vs. measurement time normalized to initial trajectory number. Left and right panels show the case without and with using an oxygen scavenging system, respectively. (**d**) Top left panel: *T*_1/2_ vs. irradiance plot for TH origami samples imaged at varying imager concentrations. Bottom left panel: Plot of mean number of detected photons per localization. Right panels: Distributions of number of photons detected per localization corresponding to concentration series imaged at *E*=30 W/cm^2^. (**e**) *T*_1/2_ vs. irradiance plots for TH origami imaged at varying temperature. (**f**) *T*_1/2_ vs. irradiance plots for TH origami imaged at varying buffer ion compositions. Error bars in b,c,d,e,f correspond to relative error determined in **Supplementary Figure 5**.

First, we examined to what extent photobleaching as a function of irradiance is a limiting factor to the TH performance. To do so, we first had to assay the average bleaching rate of single Cy3B dye molecules (remember that each imager carries a single Cy3B molecule, i.e. the average time it takes a dye molecule to photobleach should be larger than the average binding time or *k*_off_ *> k*_photobleach_) by imaging immobilized SD origami at increasing irradiances (*E*=10 W/cm^2^, 30 W/cm^2^ and 65 W/cm^2^). Faster photobleaching of Cy3B molecules was observed with increasing irradiance, as one would expect (*T*_1/2_ = 30 s, 12 s and 6 s, respectively blue decaying curve in the left panel in **Figure 3**b). Next, we imaged TH origami at the same irradiances. Despite a similar decay with increasing irradiance, we on average obtained a 26-fold increase in *T*_1/2_ compared to SD origami (orange curve in **Figure 3**b, left panel). For instance, in 30 minutes imaging of TH origami at 30 W/cm^2^, we obtained *T*_1/2_ of 365 s (> 6 minutes) compared to only 12 s for SD origami.

In order to suppress fast photobleaching, we repeated the irradiance series in the presence of the oxygen scavenging system POC (pyranose oxidase, catalase and glucose) and the triplet state quencher trolox (imaging buffer POCT, acquisition length: 30 min for SD origami and at least 60 min for TH origami). To our surprise, even for SD origami we obtained *T*_1/2_ values in the range of hundreds of seconds (blue curve in **Figure 3**b, right panel). However, for TH origami we even obtained another 5-fold increase in comparison to SD origami (orange curve). With POCT, in 60 minutes imaging at *E*=30 W/cm^2^, we obtained a *T*_1/2_ of ∼26 minutes. Repeating the measurement at *E*=10 W/cm^2^ with an extended measurement time of 3 hours we even obtained a *T*_1/2_ value of more than 1 hour (see also **Supplementary Figure 6**).

We also investigated the effect of photo-induced damage to the survival time of the TH. Analyzing the number of trajectories per frame for the TH data sets of the irradiance series clearly indicates that *k*_photodamage_ increases with higher irradiances (**Figure 3**c, left panel). In **Supplementary Figure 7** we show that *k*_photodamage_ follows a linear dependence on the irradiance. However, despite this damage occurring over time, even at *E*=65 W/cm^2^ (where SD origami had a *T*_1/2_ of 6 s) on average more than 60 % of all THs were detected in every frame over 30 minutes. The source of the damage lies in reactive oxygen species, confirming previous results^17,19^(**Figure 3**c, right panel). These can be efficiently removed using POCT maintaining a constant level of detected THs independent of the applied irradiance over the same time interval. As previously mentioned, we optimized the TH performance in a buffer compatible with live cell conditions (buffer L, T=21 °C, 40 nM imager). For future specific SPT problems, one or more of these conditions might have to be adapted. In the following, we therefore analyzed the effect of changes to imager concentration, temperature and buffer composition (salt), keeping the other parameters fixed, with respect to TH function. First, we varied the imager concentration for samples with immobilized TH origami ([imager] = 5 nM, 10 nM, 20 nM and 40 nM, at T=21°C, buffer L). As one would expect, increasing the imager concentration resulted in an increase in trajectory durations due to a higher probability of binding to an unoccupied site on the TH (**Figure 3**d, top left panel). However, increasing from 20 to 40 nM imager concentration, the increase in *T*_1/2_ is close to saturation, particularly at high irradiances. Looking at the number of photons detected per localization (**Figure 3**d, bottom left panel), we similarly observed an increase with imager concentration because multiple emitting imagers simultaneously bind the TH. The corresponding distributions of photon counts illustrates how the probability of higher order occupancies of bound imagers increases with concentration (**Figure 3**d, right panel).

Third, we highlight the influence of temperature on TH performance. Increasing the temperature from 21 °C to 23 °C (fix: buffer L, 40 nM imager concentration) had a large effect on the average trajectory duration despite this small temperature difference, as *T*_1/2_ dropped on average by ∼30 % (**Figure 3**e). The drop is due to an increased *k*_off_, which results in faster imager dissociation and thus more frequent interruptions in the fluorescence signal (**Supplementary Figure 8**).

Lastly, we emphasize the effect of ion composition (salt) onto the TH. We performed the same irradiance series for TH origami samples using two buffers with different ion compositions: buffer L (3 mM MgCl_2_ + 140 mM NaCl) and buffer B (10 mM MgCl_2_) at T=21 °C and 5 nM imager concentration. The buffer with the higher amount of Mg^2+^ ions, buffer B, resulted in longer trajectories by ∼40 % compared to buffer L due to the beneficial effect of higher MgCl_2_ concentrations to both *k*_on_ and *k*_off_ (**Figure 3**f, **Supplementary Figure 9**).

### SPT of DNA origami on Supported Lipid Membranes

Having gained a deeper understanding of the TH principle in the surface-immobilized case, we finally investigated the improvement of TH labeling compared to single-dye labeling for SPT on moving targets (origami), using 8 biotin anchors on the origami for their permanent attachment to streptavidin-biotin-functionalized supported lipid bilayers (SLB). Movement of the origami was thus governed by the lateral diffusion of the biotinylated lipids within the SLB, and the multiple lipid anchors introduced a slow two-dimensional diffusion (**Supplementary Videos 1 & 2**). A detailed description of our analysis workflow for moving particles is given in **Supplementary Figure 10**. **Figure 4**a shows the 16 longest trajectories we obtained under identical imaging conditions of both the SD origami (top) and the TH origami (bottom) ‘floating’ on the SLB. While the spatial dimensions of the trajectories confirmed the diffusivity of the system in both cases we could readily observe that even the shortest trajectory of the TH origami (> 240 s) is longer in duration than any of the trajectories we obtained for the SD origami (< 120 s). Remarkably, we were able to observe a single TH origami for more than 18 min during this measurement (for other irradiances see **Supplementary Figure 11**).

**Figure 4.**
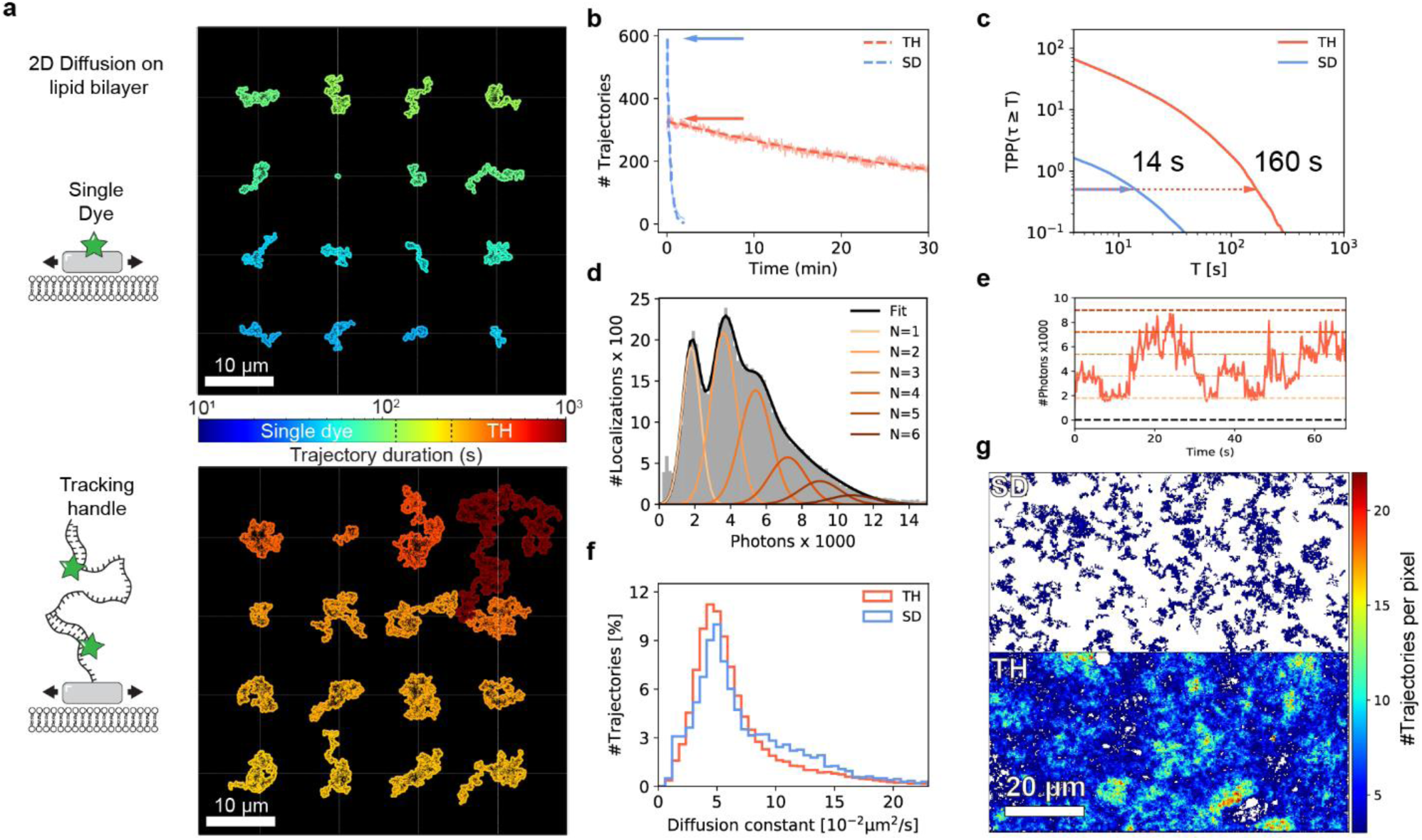
Probing 2D diffusion of DNA origami on lipid membranes. (**a**) the 16 trajectories of longest duration of both SD origami (top) and TH origami (bottom) floating on an SLB. The trajectories where shifted according to their mean position indicated by the joints of the white dashed lines and were color-coded by duration for better visibility. (**b**) Number of trajectories per frame for SD origami (solid blue) and TH origami (solid orange). The curves were fitted by an exponential decay function (dashed). The arrows indicate the initial track number (∼number of Origami) *M* within the FOV. (**c**) Average number of tracks per Origami *TPP*(*τ*_*n*_ ≥ *T*) as obtained by normalization to the initial track number *M* (arrows in b). (**d**) Photon counts per localization for the moving TH origami in the circular center of the FOV (diameter = 400 px) (grey). We applied the same fit model as in **Figure 2**f to analyze how many emitting imagers are bound to the TH (sum in black, individual colored). (**e**) Fluorescence trace of an exemplary TH trajectory revealing step like behavior and the number of currently emitting imagers bound to the TH. The dashed horizontal lines indicate the center of the 6 Gaussians obtained in (d). (**f**) Diffusion constants as obtained by linear iterative fitting of the individual MSD curves. Histograms represent the total distribution of three different samples imaged under three different irradiances. (**g**) Number of unique trajectories per 2×2 binned pixel. The TH origami allowed almost complete mapping of the SLB for (bottom half) in contrast to only sparse sampling for the SD origami (top half). White areas indicate that no trajectory passed these pixels over the complete measurement time. Scale bars, 10 *μ*m in (a), 20 *μ*m in (g).

Again, we computed the number of trajectories per frame over the course of the measurement (**Figure 4**b, solid lines) and fitted an exponential decay (dashed lines). For SD origami, we almost did not observe any emitting labels already after two minutes yielding a *T*_1/2_ value of ∼15 s (**Figure 4**b, blue line), which is in good agreement with the results obtained from the immobilized samples (compare **Figure 2**c,h). In contrast, for the TH origami, we still registered more than half of the initial trajectory number per frame at the end of the measurement duration of 30 min (half time ∼33 min, **Figure 4**b, orange line). In total, we collected ∼19,000 trajectories over the course of the measurement which represents a 20-fold (statistical) increase compared to the SD origami (only ∼900). **Supplementary Figure 12** shows an analogous analysis under different irradiances.

In SPT, it is generally not possible to unambiguously identify multiple appearances of the same mobile particle in a data set. Hence, every reappearance of a diffusing DNA origami led to a new trajectory of duration *τ*_*n*_. The total number of trajectories of a data set we defined as *N*_*tot*_. We could recover the average number of trajectories per particle 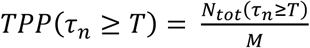 with a duration *τn* longer or equal to the time *T* by dividing the total number of trajectories *N*_*tot*_(*τ*_*n*_ ≥ *T*) by the initial trajectory number *M*. Note that the initial trajectory number (marked by arrows in **Figure 4**b) should give a good estimate for the number of particles *M* within the FOV for sufficiently low particle densities. This allowed us to directly compare trajectory durations of both immobile and mobile particles using the same metric (compare **Figure 2**c,d). To control the applicability of this approach we reanalyzed the immobilized TH origami using this procedure (see **Supplementary Figure 13**). **Figure 4**c shows *TPP*(*τ*_*n*_ ≥ *T*) for the diffusing SD origami (blue) and TH origami (red). *TPP*(*τ*_*n*_ ≥ *T*) was < 1.5 for the SD origami, confirming the validity of our approach since we ideally would expect only one trajectory per single dye before it photobleaches (compare **Figure 2**c). Furthermore, the *T*_1/2_ value of ∼12.7 s is in good agreement with the *T*_1/2_ of ∼12 s of the immobilized SD origami (compare **Figure 2**c. See **Supplementary Figure 12** for additional irradiances). The TH origami yielded a *T*_1/2_ of ∼160 s prolonging the on average trajectory duration compared to the single dye by a factor > 12x. However, if we compare this value to the *T*_1/2_ = 365 s of the immobilized samples imaged under identical conditions, we notice a shortening by more than a factor of two. We reason that the reduced trajectory durations *T*_1/2_ of the diffusing TH origami is due to the limited capability of our (nearest-neighbor based) linking algorithm at the used particle density (see **Supplementary Figure 14**).

Analogous to **Figure 2**f we extracted and analyzed the photon counts per localization of the diffusing TH origami (**Figure 4**d). Similar to the immobilized case, we were able to observe up to six discrete photon levels indicating a multitude of imagers bound to the TH (for the photon count histogram of the single dye origami refer to **Supplementary Figure 15**). Further evidence is given by the good visual alignment of the centers of the Gaussian fits with the fluorescence trace of an exemplary trajectory shown in **Figure 4**e.

Next we investigated the diffusion properties of both the SD origami and the TH origami on the SLB, obtaining the diffusion constants by linear (iterative) fitting of the individual mean square displacement (MSD) curves (**Figure 4**f). The histogram presents the total distribution of three different samples imaged under three different iraddiances of all trajectories containing more than 20 localizations giving a total of ∼61.000 trajectories for the TH origami and ∼9.000 trajectories for the SD origami (see **Supplementary Figure 12** for individual irradiances). The resulting distributions agree well with each other, indicating that the diffusion properties are not altered by the TH.

Finally, following previous ideas of to create spatially resolved maps of single-molecule motions^21,22^, we divided the complete FOV into binned pixels of size 2×2. For each binned pixel we counted the number of unique trajectories passing through it during the entire measurement (*i.e.* a unique trajectory having at least one localization within the pixel boundaries increases its count by one). The resulting maps for half the FOVs are shown in **Figure 4**g for both the SD origami (top half) and the TH origami (bottom half). Both the high number of registered trajectories per single TH origami and the long duration of the individual trajectories allowed an almost complete mapping of the SLB with only ∼350 origami present in the FOV at the start of the measurement (arrow in **Figure 4**b). Whereas the map allowed to identify either inaccessible areas or areas without membrane coverage (white) in the case of the TH origami the trajectories of the SD origami could only cover the membrane in parts not allowing any conclusions about its morphology (despite the elevated particle density, **Figure 4**b).

### The power of long particle trajectories

One of the most prominent features of the TH is the generation of significantly longer particle trajectories when compared to single-dye labeling, reaching durations of up to tens of minutes. We hence want to clearly point out how the precision of the experimental outcome is influenced by the duration of the observed particle trajectories. As previously mentioned, we obtained the diffusion coefficient *D* by linear (iterative) fitting of the individual MSD curves (see **Supplementary Figure 10**). In this case, numerous theoretical works describe a loss in precision in the obtained diffusion coefficient with decreasing trajectory duration^23–25^. In the following we want to assess this behavior experimentally based on the data of the floating TH origami.

We divided all trajectories exceeding 120 s (600 frames) in duration into subtrajectories of 10 s (50 frames), as illustrated in **Figure 5**a. After this, we applied the same MSD fitting algorithm to both the complete trajectory and to all of its subtrajectories. Each trajectory hence yielded one diffusion coefficient *D* (using its full length) and multiple diffusion coefficients *D*_*sub*_ for each of its subtrajectories. **Figure 5**a shows an exemplary trajectory of a TH origami and the corresponding subtrajectories color coded by the obtained diffusion coefficients *D*_*sub*_.

**Figure 5.**
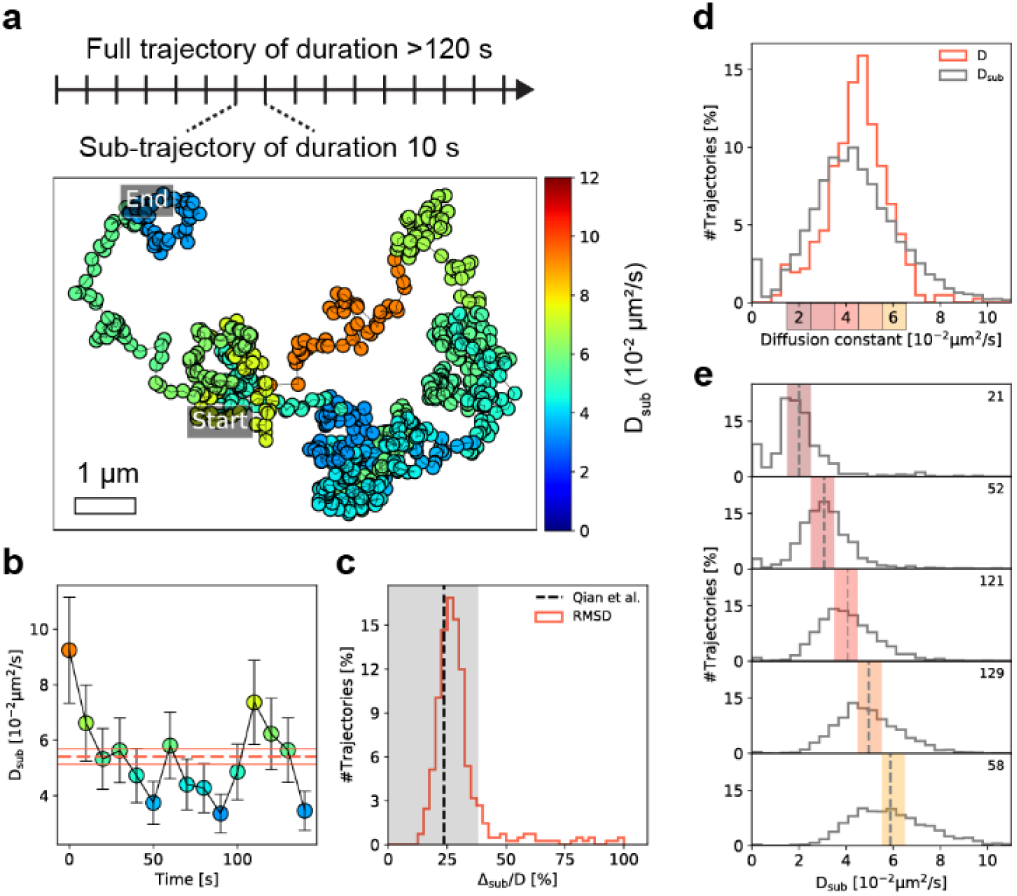
Subtrajectory analysis of TH origami diffusion on SLB. Trajectories exceeding 120 s in duration were divided into subtrajectories of duration 10 s and analyzed by the same MSD fitting procedure. (**a**) In the exemplary trajectory each localization is color-coded according to the obtained diffusion constant *D*_*sub*_ of the corresponding subtrajectory. (**b**) Time dependent scatter of *D*_*sub*_ (see colorbar in a) around the diffusion constant *D* (red solid, error: red dashed) as obtained from analysis of the full trajectory. Errors for *D*_*sub*_ and *D* are calculated according to Eq. 1. (**c**) RMSD distribution of *D*_*sub*_ to *D* of all trajectories exceeding 120 s (red). RMSD was normalized to *D* and should hence be close to the theoretical limit (black dashed) if the TH origami are subject to a time-invariant Brownian motion. Grey area indicates deviation of less than 60% to the theoretical limit. (**d**) Total distribution of *D* and the corresponding subtrajectory diffusion constants *D*_*sub*_. (**e**) We selected five subsets of trajectories yielding a value of *D* within the ranges 1.5 - 2.5, …, 5.5 – 6.5 *μ*m^2^/s (colored boxes in d) and plotted the corresponding *D*_*sub*_ distribution. The mean value of *D*_*sub*_ (grey dashed) agrees well with the selected central *D*. The top right number indicates how many trajectories were part of the selected subsets in *D*.

**Figure 5**b illustrates how *D*_*sub*_ randomly scatters around *D* as obtained from MSD analysis of the full trajectory (red dashed, error: red solid). The errors indicate the theoretical standard deviation of the diffusion coefficient for Brownian motion derived by Qian et al.^23^:

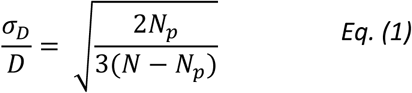

with N being the total duration of the trajectory in frames (*N* = 600, subtrajectories: *N* = 50) and *N*_*p*_ being the maximum lag time (in frames) up to which the MSD curve was fitted (see **Supplementary Figure 10**). In our case *N*_*p*_ was chosen automatically during the iterative fitting process^24^ resulting in a median value of *N*_*p*_ = 3 on average.

Naturally, the question arises: is the scatter of *D*_*sub*_ only governed by the random nature of the motion according to Eq. (1) or does an individual TH origami undergo a change in its diffusive properties during the observation time? To test this we calculated the root-mean-square deviation (RMSD) 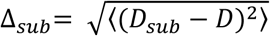 of *D*_*sub*_ to *D* for each trajectory^26^. **Figure 5**c shows the distribution of the thus obtained RMSD normalized to the diffusion coefficient *D* of the full trajectory. If the movement of the TH origami is indeed governed by the same 2D Brownian motion with a diffusion coefficient *D* at any time the RMSD Δ_*sub*_ should correspond to the standard deviation *σ*_*D*_ (for *N* = 50) and should hence yield a value close to theoretically achievable limit given by Eq. (1) (black dashed line). **Figure 5**c shows that around 86 % of the analyzed trajectories do not deviate from the expected statistical uncertainty by more than 60 % (grey area) suggesting that the TH origami are indeed subject to a time-invariant Brownian motion on the SLB. This statement is further supported by division into longer (*N* = 100) or shorter (*N* = 25) subtrajectories (see **Supplementary Figure 16**).

**Figure 5**d shows the distribution of diffusion coefficients *D* of all analyzed trajectories (red) and the corresponding diffusion coefficients *D*_*sub*_ of its 10-second subtrajectories (grey). As expected, we observed a broadening of the *D*_*sub*_ distribution by a factor of 1.5x with respect to *D* due to the larger statistical uncertainty with decreasing trajectory duration. **Supplementary Figure 17** further illustrates the influence of the subtrajectory length (*N* = 100, 25) on the broadening in the resulting *D*_*sub*_ distribution.

Even the full TH origami trajectories showed a relatively broad range of diffusion constants ranging from 2-6 *μ*m^2^/s (see **Figure 5**d). We reason that this is caused by varying numbers of biotinylated staple strands per TH origami due to a limited incorporation efficiency^27^.

To highlight the full extent of the importance of the trajectory durations decoupled from the variability of this system, we selected trajectories which yielded a similar diffusion coefficient as indicated by the colored ranges on the x-axis in **Figure 5**d (1.5 - 2.5, …, 5.5 – 6.5 *μ*m^2^/s). **Figure 5**e shows the corresponding distributions resulting from MSD analysis of the 10-second subtrajectories (grey) and the respective means (grey dashed lines). The legends states from how many full trajectories the subtrajectories originated in each range. The means of *D*_*sub*_ are located at the center of the selected ranges in *D* for all cases. However, the relative standard deviation of *D*_*sub*_ increased with respect to *D* by a factor of 4.8x on average. Calculation of the lower limit of the relative standard deviation according to Eq. (1) yields an value of ∼5 % for the full trajectory and an value of ∼24 % for its subtrajectories. Hence, the theoretically expected broadening of 24/5 = 4.8x matches well with our experimental observations confirming a time-invariant 2D Brownian motion of the TH origami on the SLB (see **Supplementary Figure 17** for other subtrajectory durations).

## Discussion

Here we presented our new labeling strategy for fluorescence-based SPT, using a 1:1 functionalization with a DNA-based TH and exploiting DNA-mediated fluorophore exchange to increase observation times. By largely decoupling the trajectory duration from the photon budget of single dye molecules, we showed that our TH allows observation of target particles from minutes to hours, depending on the experimental conditions. However, this comes at a cost: the TH needs to be accessible to freely-diffusing imagers in solution, which, typically for DNA-PAINT, currently limits TH experiments to selective plane illumination schemes due to the high fluorescence background. Very recent development on fluorogenic imager design could possible overcome this characteristic DNA-PAINT constraint^28^. Additionally, negative control experiments without THs but with imagers present in the solution are required to account for the possibility of unspecific imager binding.

At the example of 2D diffusion on SLBs, we showed numerous advantages using the TH compared to single-dye labeling. The large number of trajectories led to a coverage of the whole FOV which allowed to map the whole accessible area with an actual low number of particles. Even for moving THs, the number of currently bound imagers can be recovered from step-like intensity fluctuations in the fluorescence trace, which provides the potential for intensity barcoding^29^ and multiplexing^30^ in the future beyond the use of orthogonal DNA sequences. The ability to divide long trajectories into subtrajectories paves way for a robust quantitative analysis of the underlying dynamic^26,31,32^. While we here confirmed a time-invariant 2D Brownian diffusion as the driving force and experimentally verified the widely-accepted theory in the field, this concept can potentially be translated to anomalous diffusion behavior such as stop-and-go motion. Likewise, long trajectories are essential to unambiguously identify regions of confined motion and molecular interactions in SPT experiments, which are often paramount to biological function. Our detailed analysis of the key parameters of DNA-PAINT based SPT will allow experimenters to adapt the TH to their needs, ranging from *in vitro* applications to potentially tracking receptors in the plasma membrane of living cells. We believe that due to its variability and simple implementation the TH will become a valuable tool for studying dynamic processes at the single molecule level.

## Acknowledgements

We thank Patrick Schueler, Beatrice Ramm, Tamara Heermann, Henri Franquelim, Sigrid Bauer, and Katharina Nakel for their support and helpful discussions.

## Data availability statement

All raw data is available upon reasonable request from the authors.

