## Supplementary Information for "Tracking Single Particles for Hours via Continuous DNA-mediated Fluorophore Exchange"



| | |
|---|---|
| **Supplementary Methods** | |
| **Supplementary Figure 1** | **Size comparison between SPT labels** |
| **Supplementary Figure 2** | **Data analysis workflow - immobilized** |
| **Supplementary Figure 3** | **Filter - immobilized** |
| **Supplementary Figure 4** | **Single TH trace photon histogram** |
| **Supplementary Figure 5** | **Error estimation for $T_{1/2}$ results** |
| **Supplementary Figure 6** | **3 hour tracking with POCT** |
| **Supplementary Figure 7** | **$k_{photodamage}$ vs irradiance** |
| **Supplementary Figure 8** | **Temperature effect on DNA hybridization rates** |
| **Supplementary Figure 9** | **Ion concentration effect on DNA hybridization rates** |
| **Supplementary Figure 10** | **Data analysis workflow - mobile** |
| **Supplementary Figure 11** | **Longest trajectories at varying irradiances** |
| **Supplementary Figure 12** | **Tracks, TPP and diffusion constant for varying irradiances** |
| **Supplementary Figure 13** | **Mobile analysis control with immobilized data set** |
| **Supplementary Figure 14** | **Linking and particle density** |
| **Supplementary Figure 15** | **Photons counts - SD vs TH mobile** |
| **Supplementary Figure 16** | **RMSD histograms for N=25 & N=100** |
| **Supplementary Figure 17** | **D vs. $D_{sub}$ for N=25 & N=100** |
| **Supplementary Table 1** | **Imaging parameters** |
| **Supplementary Table 2** | **Used DNA-PAINT sequences** |

#### Supplementary methods

##### Materials

Unmodified, dye-labeled, and biotinylated DNA oligonucleotides were purchased from MWG Eurofins. DNA scaffold strands were purchased from Tilibit (cat. p7249, identical to M13mp18). Streptavidin was purchased from Thermo Fisher (cat. S-888). BSA-Biotin was obtained from Sigma-Aldrich (cat. A8549). All lipids were purchased from Avanti Polar Lipids. Glass slides were ordered from Thermo Fisher (cat. 10756991) and coverslips were purchased from Marienfeld (cat. 0107032). Freeze 'N Squeeze columns were ordered from Bio-Rad (cat. 732-6165). Tris 1M pH 8.0 (cat. AM9856), EDTA 0.5M pH 8.0 (cat. AM9261), Magnesium 1M (cat. AM9530G) and Sodium Chloride 5M (cat. AM9759) were ordered from Ambion. Ultrapure water (cat. 10977-035) was purchased from Thermo Fisher Scientific. Tween-20 (cat. P9416-50ML), Glycerol (cat. 65516-500ml), (+-)-6-Hydroxy-2,5,7,8-tetra-methylchromane-2-carboxylic acid (Trolox) (cat. 238813-5G), pyranose oxidase (PO, cat. P4234) and catalase (C, cat. C40) were purchased from Sigma-Aldrich. Two-component epoxy glue (cat. 886519 - 62) was purchased from Conrad Electronic SE.

##### Buffers

Seven buffers were used for sample preparation and imaging: Buffer A (10 mM Tris-HCl pH 7.5, 100 mM NaCl); Buffer B (5 mM Tris-HCl pH 8.0, 10 mM $MgCl_2$, 1 mM EDTA); Buffer L (20 mM HEPES pH 7.6, 140 mM NaCl, 3 mM MgCl2); Buffer POCT (5 mM Tris-HCl pH 8.0, 10 mM $MgCl_2$, 1 mM EDTA, incubated 1 h prior to measurement with 1x PO, 1x C, 0.8 % Glucose and 1x Trolox as previously described[1]); 10x folding buffer (100 mM Tris,10 mM EDTA pH 8.0, 125 mM MgCl2); Buffer M (25 mM Tris HCl pH 7.5, 150 mM KCl, 5 mM MgCl2); SLB buffer (25 mM Tris HCl pH 7.5, 150 mM KCl).

##### DNA origami design and assembly and purification

DNA origami structures were designed using the design module of Picasso[2]. Our DNA origami design is identical to the one used in previous work[3], i.e. of flat rectangular geometry with just a single extension on the top side (at position 2B07 of Picasso Design). At this position, we as the label either incorporated the TH or a Cy3B molecule permanently attached to a short T-spacer, or a single DNA-PAINT docking strand (1DS) (see **Supplementary Table 2** for sequences). On the bottom side, 8 biotinylated extensions were incorporated for surface immobilization/SLB binding. Folding of structures was performed using the following components: single-stranded DNA scaffold (0.01 µM), core staples (0.1 µM), biotin staples (1 µM for SD origami and 0.01 µM for TH origami and 1DS origami), TH/SD/1DS strands (1 µM), 1× folding buffer in a total of 50 µl for each sample. Annealing was done by cooling the mixture from 80 to 25 °C in 3 h in a thermocycler. TH origami and 1DS were not purified after folding. SD origami were purified using PEG precipitation[4].

**Sample preparation**

Surface-immobilized DNA origami

DNA origami samples were prepared as described before[2]. A glass slide was glued onto a coverslip with the help of double-sided tape (Scotch, cat. no. 665D) to form a flow chamber with inner volume of ~20 µl. First, 20 µl of biotin-labeled bovine albumin (1 mg/ml, dissolved in buffer A) was flushed into the chamber and incubated for 3 min. The chamber was then washed with 40 µl of buffer A. 20 µl of streptavidin (0.5 mg/ml, dissolved in buffer A) was then flushed through the chamber and incubated for 3 min. After washing with 40 µl of buffer A and subsequently with 40 µl of buffer B, 20 µl of biotin-labeled DNA origami (dilution from DNA origami stock dependent on origami yield after gel purification. Adjusted for each origami species individually to obtain a sparse DNA origami surface density. Starting dilution ~1:200) were flushed into the chamber and incubated for 10 min. The chamber was washed with 80 µl of imaging buffer (L/B/POCT) to remove unbound DNA origami. SD origami samples were sealed with two-component epoxy glue before imaging. For TH and 1DS origami samples, 40 µl of the imager solution was flushed into the chamber before sealing.

DNA origami diffusing on supported lipid bilayers

A glass slide was rubbed and rinsed with EtOH and dd$H_2O$ and subsequently plasma cleaned using a plasma cleaner (Zepto, Diener Electronic, Germany) for 40 s at 50% power and 0.3 mbar with oxygen as process gas. The glass slide was glued onto a coverslip with the help of double-sided tape (Scotch, cat. no. 665D) to form a flow chamber with inner volume of ~20 µl. Small unilamellar vesicles (SUVs) were prepared at a concentration of 4 mg/ml in buffer M from a lipid composition of 99 mol % DOPC/1 mol % Biotinyl-CAP-PE. Lipids dissolved in chloroform were dried under a stream of nitrogen. Vials were placed in a desiccator for 30 min to remove residual chloroform. After lipids were rehydrated in 200 µl of buffer M, the vials were placed in a sonicator bath to generate SUVs until the solution appeared transparent (storage of SUV solution aliquots possible at -30 °C for several weeks. After thawing of an aliquot it was placed in the sonication bath for 30 min). SUV solution was diluted to 0.5 mg/ml in buffer M. 20 µl of SUV dilution was flushed into the chamber and incubated for 3 min. The chamber was then washed with 5x80 µl of SLB buffer to remove excess vesicles and 1x80 µl with buffer B. 20 µl of streptavidin (0.5 mg/ml, dissolved in buffer A) was then flushed through the chamber and incubated for 5 min. After washing with 80 µl of buffer B, 20 µl of biotin-labeled DNA origami was flushed into the chamber and incubated for 3 min (origami dilution after folding ~1:1,000). Excess DNA origami were washed with 80 µl of buffer L. SD origami samples were sealed with two-component epoxy glue before imaging. For TH origami samples 40 µl of the imager solution were flushed into the chamber before sealing.

**Super-resolution microscopy setup**

Fluorescence imaging was carried out on an inverted custom-built microscope (see supplementary references[3,5] for detailed sketches) in an objective-type TIRF configuration with an oil-immersion objective (Olympus UAPON, 100×, NA 1.49). One laser was used for excitation: 561 nm (1 W, DPSS-system, MPB). The laser power was adjusted via polarization rotation using a half-wave plate (Thorlabs, WPH05M-561) before passing a polarizing beam-splitter cube (Thorlabs, PBS101). The laser light was coupled into a single-mode polarization-maintaining fiber (Thorlabs, P3-488PM-FC-2) using an aspheric lens (Thorlabs,

C610TME-A) in order to spatially clean the beam-profile. Using a zero-order half wave plate (Thorlabs, WPH05M-561) the coupling polarization into the fiber was adjusted. The laser light was re-collimated after the fiber using an achromatic doublet lens (Thorlabs, AC254-050-A-ML) resulting in a collimated FWHM beam diameter of ~6 mm. The Gaussian laser beam profile was transformed into a collimated flat-top profile using a refractive beam shaping device (AdlOptica, piShaper 6_6_VIS). The laser beam diameter was magnified by a factor of 2.5 using a custom-built telescope (Thorlabs, AC254-030-A-ML and Thorlabs, AC508-075-A-ML). The laser light was coupled into the microscope objective using an achromatic doublet lens (Thorlabs, AC508-180-A-ML) and a dichroic beam splitter (AHF, F68-785). Fluorescence light was spectrally filtered with a laser notch filter (AHF, F40-072) and a bandpass filter (AHF Analysentechnik, 605/64) and imaged on a sCMOS camera (Andor, Zyla 4.2) using a tube lens without further magnification (Thorlabs, TTL180-A) resulting in an effective pixel size of 130 nm (after 2 × 2 binning). Microscopy samples were mounted into a closed water-based temperature chamber (Okolab, H101-CRYO-BL) on an x-y-z stage (ASI, S31121010FT and ASI, FTP2050) that was used for focusing with the microscope objective being at fixed position. The temperature of the objective was actively controlled using the same water cycle as the temperature chamber. Focus stabilization was achieved via the CRISP autofocus system (ASI @ 850 nm) in a feedback loop with a piezo actuator (Piezoconcept, Z-INSERT100) moving the sample. The CRISP was coupled into the excitation path of the microscope using a long pass dichroic mirror (Thorlabs, DMLP650L). Our custom TIRF setup was used for all presented data.

**Imaging conditions**

All fluorescence microscopy data was recorded with our sCMOS camera (2048 × 2048 pixels, pixel size: 6.5 µm). The camera was operated with the open source acquisition software µManager[6] at 2x2 binning and cropped to the center 700 × 700 pixel FOV. The exposure time was set to 200 ms, the read out rate to 200 MHz and the dynamic range to 16 bit. The laser power was set to a homogeneous (flat-top profile, see setup description) with a measured excitation beam diameter of d $\sim$ 130 $\mu$m[5]. We performed all experiments at laser excitation powers $P$ of 1.4 mW, 4.1 mW and 8.9 mW (as measured at the fiber exit port). We hence calculated an (upper limit) homogenous excitation irradiance $E$ at the sample space of 10 W/cm$^2$, 30 W/cm$^2$ and 65 W/cm$^2$ by using $E = 4P/\pi d^2$. SD origami samples were imaged repeatedly at 3x field of views (FOVs) for increased statistics, TH origami samples only at 1x FOV. For detailed imaging parameters specific to the data presented in all main and supplementary figures refer to **Supplementary Table 1**.

**Image processing & single particle tracking analysis**

We employed two custom written python packages for our analysis https://github.com/schwille-paint/picasso_addon (*picasso_addon*) and https://github.com/schwille-paint/SPT (*spt*). These packages are based on https://github.com/jungmannlab/picasso (*picasso*) for localization of raw images and on https://soft-matter.github.io/trackpy/v0.4.2/ (*trackpy*) for linking of localizations into particle trajectories. The custom packages *picasso_addon* and *spt* offer a complete single particle tracking analysis workflow for both mobile and immobilized particles. Refer to **Supplementary Figure 2, Supplementary Figure 3** and **Supplementary Figure 10** for a detailed step-by-step guide through all processing steps of immobilized and diffusing DNA origami data, respectively. Detailed information about the *picasso_addon* and *spt* API can be found on https://picasso-addon.readthedocs.io/en/latest/index.html and https://spt.readthedocs.io/en/latest/index.html.

#### Supplementary Figures

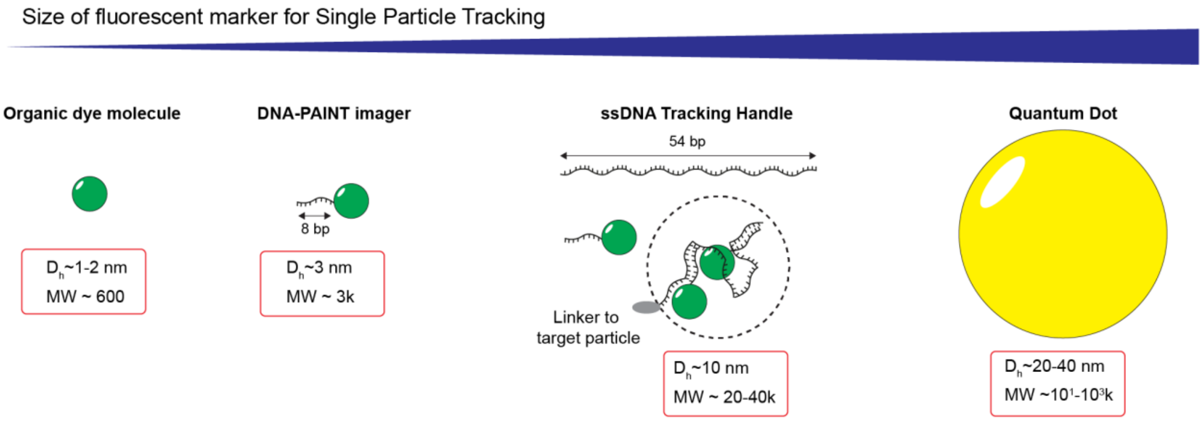


**Supplementary Figure 1. Size comparison between SPT labels.** True to scale estimation of the size (assuming a spherical shape with hydrodynamic radius $D_h$) and the molecular weight (MW) of an organic dye, a DNA-PAINT imager, the TH and a quantum dot. The $D_h$ estimation of the TH takes into account the flexible nature of ssDNA assuming coiling due to unbound regions. The MW estimation of the TH assumes on average 3x bound imagers.

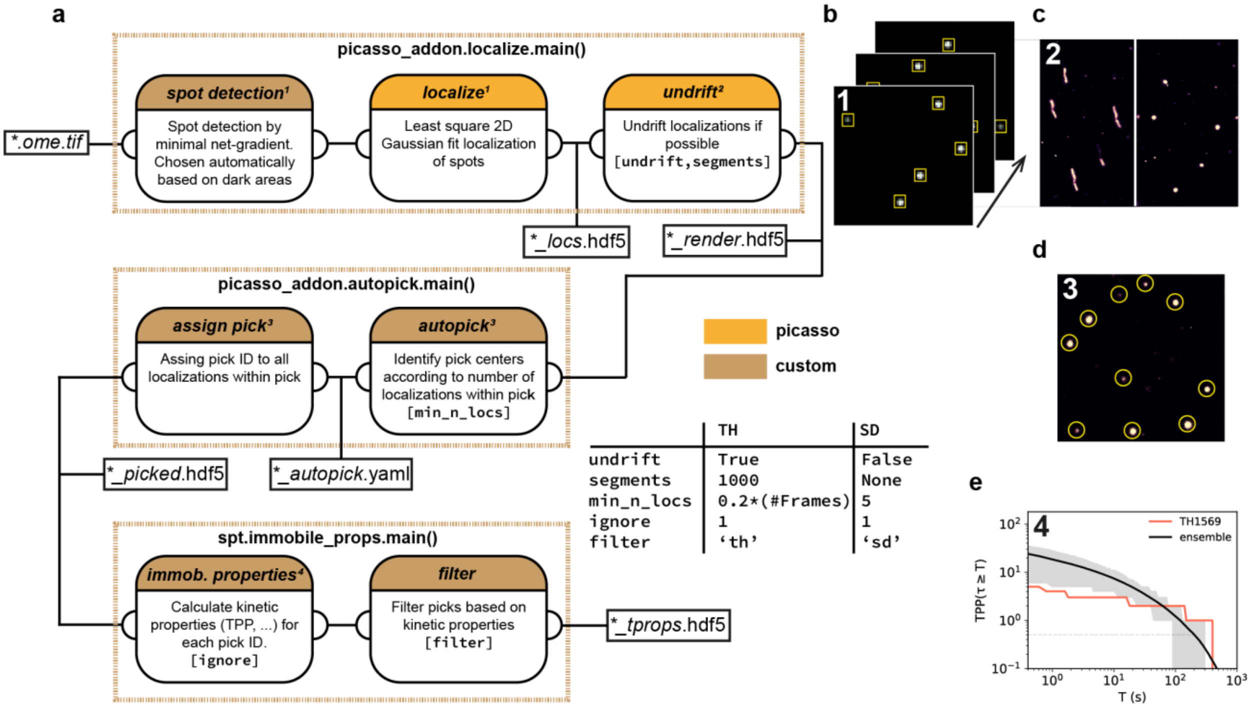


**Supplementary Figure 2. Data analysis workflow - immobilized.** (**a**) Data analysis workflow from raw movies (*.ome.tif) to final result (*_tprops.hdf5). Dashed boxes indicate employed main functions - e.g. picasso_addon.localize.main() - of the *picasso_addon* and *spt* python package. Rounded boxes illustrate provided functionalities of each main function. The header color code indicates from which python package the functionality was adapted (e.g. *picasso* or *custom* for extended functionalities within *picasso_addon* or *spt*). The text within the rounded boxes gives a short description of the provided functionality and parameters (brackets) for execution of the main function. The table summarizes all parameters used for evaluation of immobilized SD or TH experiments. The small boxes branching of the main flow represent which files are saved during execution. Please visit the links provided in the section "**Image processing & single particle tracking analysis**" for further information. (**b**) Illustration of spot detection (boxes) and localization of individual emitters in raw images. (**c**) Illustration of image correlation based undrifting of the rendered localization lists (paramters: undrift, segments). (**d**) Illustration of localization cluster detection based on number of localizations within the localization cluster (parameter: min_n_locs). We follow the Picasso nomenclature referring to a detected localization cluster as 'pick' with a unique pick ID. (**e**) Final result is obtained by calculating kinetic properties for each pick (e.g. TPP, number of localizations, photon counts etc.) by employing spt.immobile_props.main() (parameters: ignore, filter). For a detailed description of our final filtering procedure for each pick please refer to **Supplementary Figure 3.**

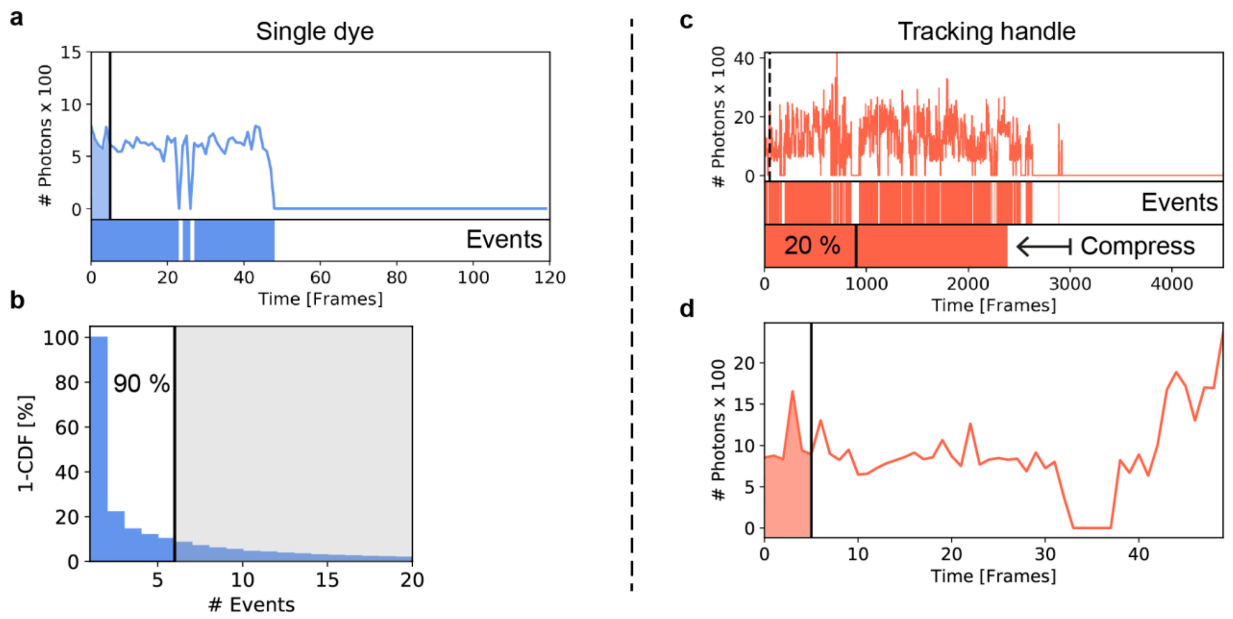


**Supplementary Figure 3. Filter - immobilized.** (**a**) The filtering procedure (see **Supplementary Figure 2**) for immobilized SD origami is illustrated for an exemplary fluorescence trace (blue line). Valid picks (i.e. picks passing the filter criteria) yielded at least one localization within the first 5 frames (black line) of the measurement (blue area below trace). The bar below the fluorescence trace indicates the number of events, i.e. a continuous and uninterrupted fluorescence signal. (**b**) Second, all picks exceeding the 90% percentile of the number of events distribution of all picks are disregarded, i.e. all fluorescence traces consisting of more than 5 events in this case. (**c**) For TH origami (illustration analogous to a) we divided the total number of localizations within a fluorescence trace by the measurement duration in frames (occupation) that can be visualized as a compression of the overall event durations. Only picks with an occupation of more than 20 % (black line) were used for further analysis. The black dashed line indicates a zoom into the fluorescence signal shown in (**d**). Analogues to (a), we additionally only considered TH origami picks yielding at least one localization within the first 5 frames (black line) of the measurement (red area below trace).

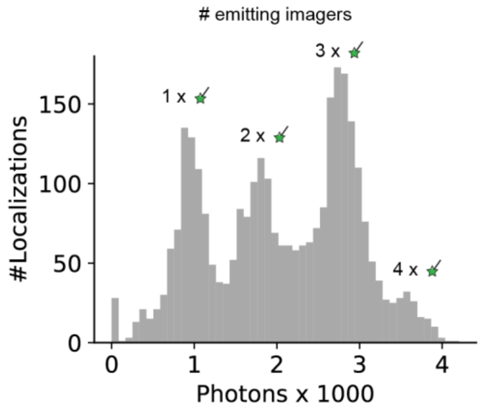


**Supplementary Figure 4. Photon count histogram individual TH trace.** The histogram corresponds to trace TH1569 displayed in **Figure 2**b. For individual traces, the equidistant peaks revealing the number of currently bound and emitting imagers (here 1-4x imagers) are more clearly separable than in the ensemble histogram from all TH origami (compare **Figure 2**f). Note that the bin at 0 displays the number of dark frames, in which no localization was detected from the TH origami (here 28 frames without localization. Remember that we ignored interruptions of just a single frame, i.e. here only interruptions ≥ 2 consecutive frames are considered).

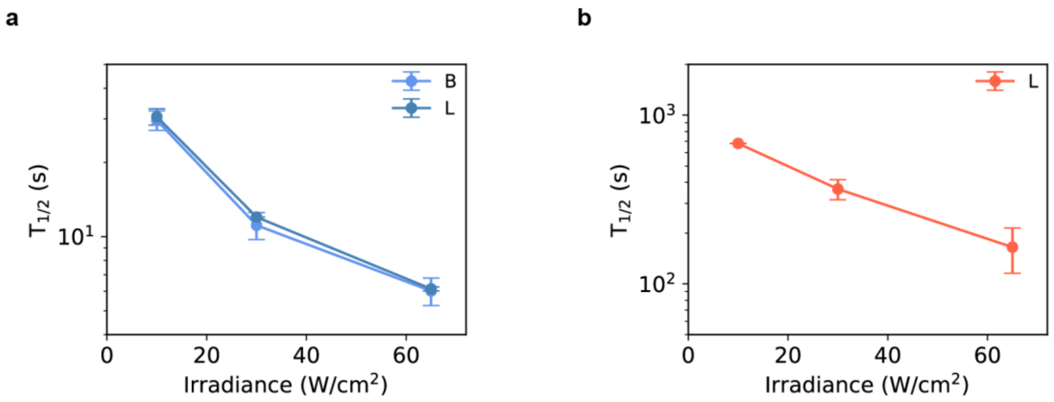


**Supplementary Figure 5. Error estimation for $T_{1/2}$ results.** (**a**) Repeated irradiance series for fresh samples of SD origami in buffer B (light blue; 4x samples; each sample imaged at 3x field of views (FOVs)) and buffer L (dark blue; 1x samples, 3x FOVs). The plots display the mean $T_{1/2}$ values averaged over all repeats per buffer condition. Error bars correspond to the standard deviation (std). The largest relative error (i.e. std/mean) was ~8 %, which used to display error bars to the SD origami results shown in **Figure 3**b. It should also be noted that the buffer ion composition did not influence the photobleaching behavior of Cy3B, since both irradiance series yielded almost the same results. (**b**) Repeated irradiance series for fresh samples of TH origami under the conditions buffer L, T=21°C and [imager]=40 nM (2x samples). The plots display the mean $T_{1/2}$ values averaged over the two repeats. Error bars correspond to the standard deviation (std). The largest relative error was ~30 %, which was used to display error bars to the TH origami results shown in **Figure 3 (**b,d,e,f).

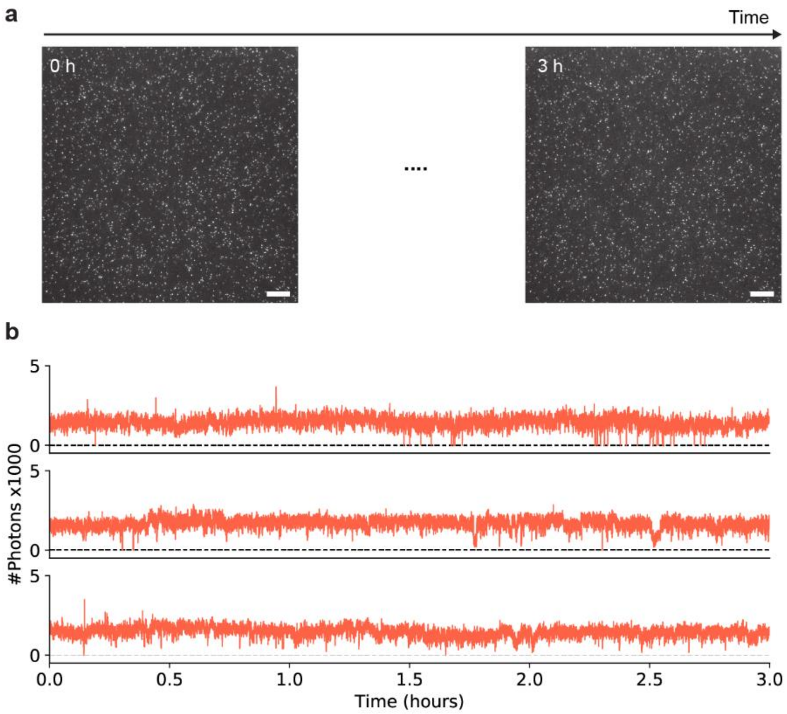


**Supplementary Figure 6. Extended 3-hour measurement using POCT.** (**a**) First and last frame of the 3h TIRFM acquisition of static TH origami still emitting after the course of the measurement. Parameters: buffer POCT, [imager]=40 nM, T=21 °C, $E$=10 W/cm$^2$. (**b**) Three exemplary fluorescence traces exhibiting continuous intensity fluctuations over 3 h. The combination POC and trolox suppresses photobleaching and photodamage to the TH. However, still short interruptions due to the stochastic nature of DNA association and dissociation occur. Scale bars, 10 $\mu$m in (a).

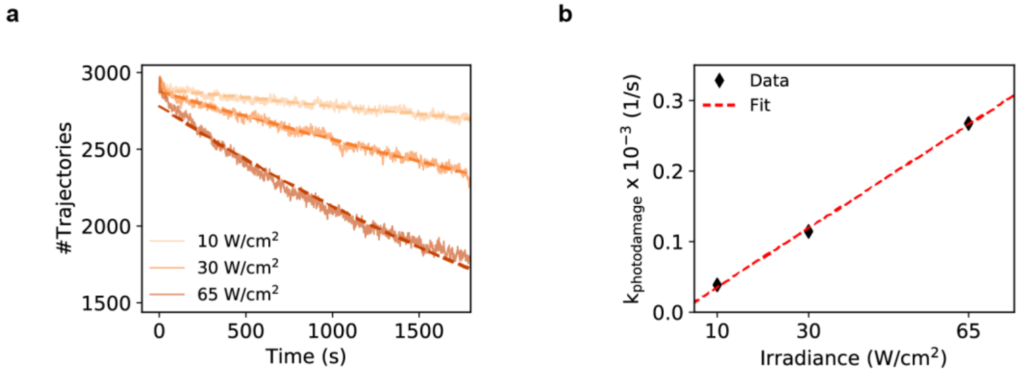


**Supplementary Figure 7. $k_{\text{photodamage}}$ vs irradiance.** (**a**) Number of trajectories per frame vs. measurement time normalized to initial trajectory number (transparent fluctuating curves) from DNA origami samples imaged at varying irradiances, as shown in left panel in **Figure 3**c. An exponential model $f(x) = ae^{-x/\tau_{photodamage}}$ was fitted to the data (dashed curves), where $\tau_{\text{photodamage}}$ denotes the characteristic decay constant over which photodamage occurs and $a$ the initial number of trajectories. (**b**) The inverse of $\tau_{\text{photodamage}}$ yielded the rate $k_{\text{photodamage}} = 1/\tau_{\text{photodamage}}$ for each irradiance. Plotting $k_{\text{photodamage}}$ vs. irradiance was well described by a line fit (red dashed line) confirming a linear relation of the two quantities over the here applied irradiance range.

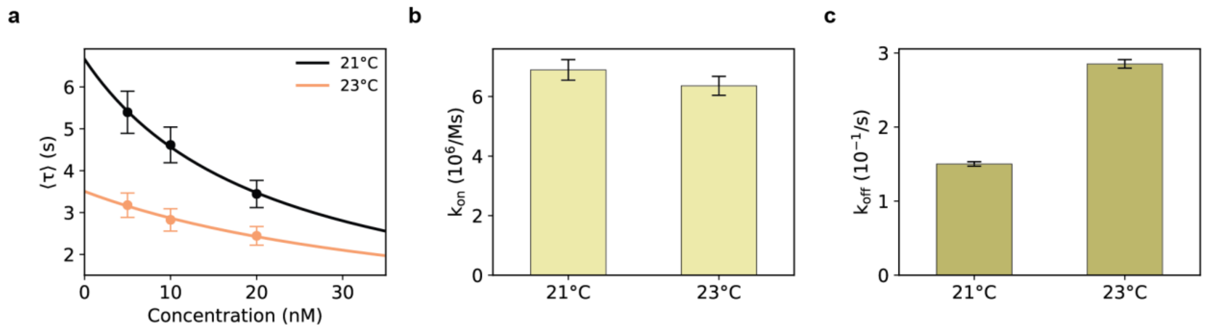


**Supplementary Figure 8. Temperature effect on DNA hybridization rates.** (**a**) As previously described[3], we performed localization-based Fluorescence Correlation Spectroscopy (lbFCS) measurements on DNA origami labeled with just a single docking strand fully complementary to the imager sequence (see **Supplementary Table 2**). By performing an imager concentration series ([imager]=5 nM, 10 nM and 20 nM) lbFCS allows to precisely measure the DNA hybridization rates $k_{on}$ and $k_{off}$[3]. Here, we conducted the imager concentration series in buffer B at T=21 °C and T=23 °C. Fitting eq. 2 in ref. 3 to the mean characteristic decay constant ⟨τ⟩ plotted vs. the imager concentration yielded $k_{on}$ and $k_{off}$. (**b**) $k_{on}$ results from the two fits in (a) indicating only minor changes due to the temperature variation. (**c**) $k_{off}$ results from the two fits in (a) highlighting the drastic temperature effect on the dissociation rate. $k_{off}$ increased almost by a factor of x2 when increasing the temperature from 21 °C to 23 °C. Error bars correspond to standard deviation.

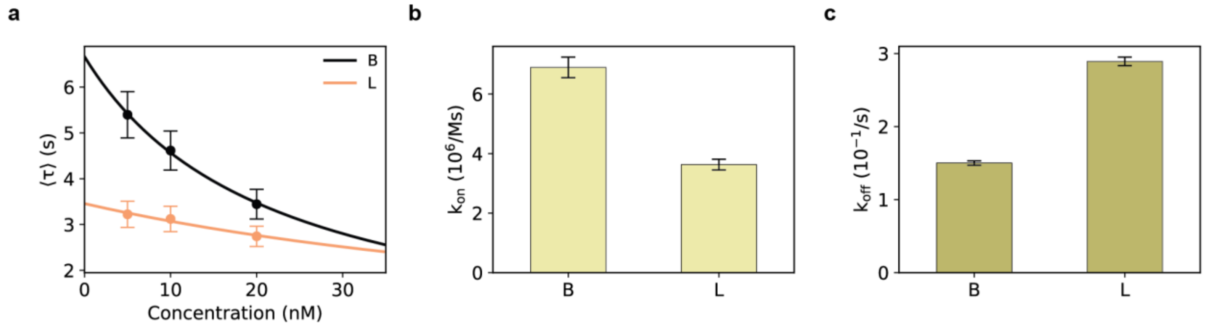


**Supplementary Figure 9. Ion concentration effect on DNA hybridization rates.** (**a**) Analogously to **Supplementary Figure 8**, we repeated the same lbFCS imager concentration series on single-docking-strand origami samples using buffer B and buffer L (at T=21 °C). (**b**) $k_{on}$ results from the two fits in (a) exhibiting a decreased association rate due to a lower $MgCl_2$ concentration in buffer L. (**c**) $k_{off}$ results from the two fits in (a). In contrast to $k_{on}$, $k_{off}$ increased nearly by a factor of 2x using buffer L compared to buffer B. Error bars correspond to standard deviation.

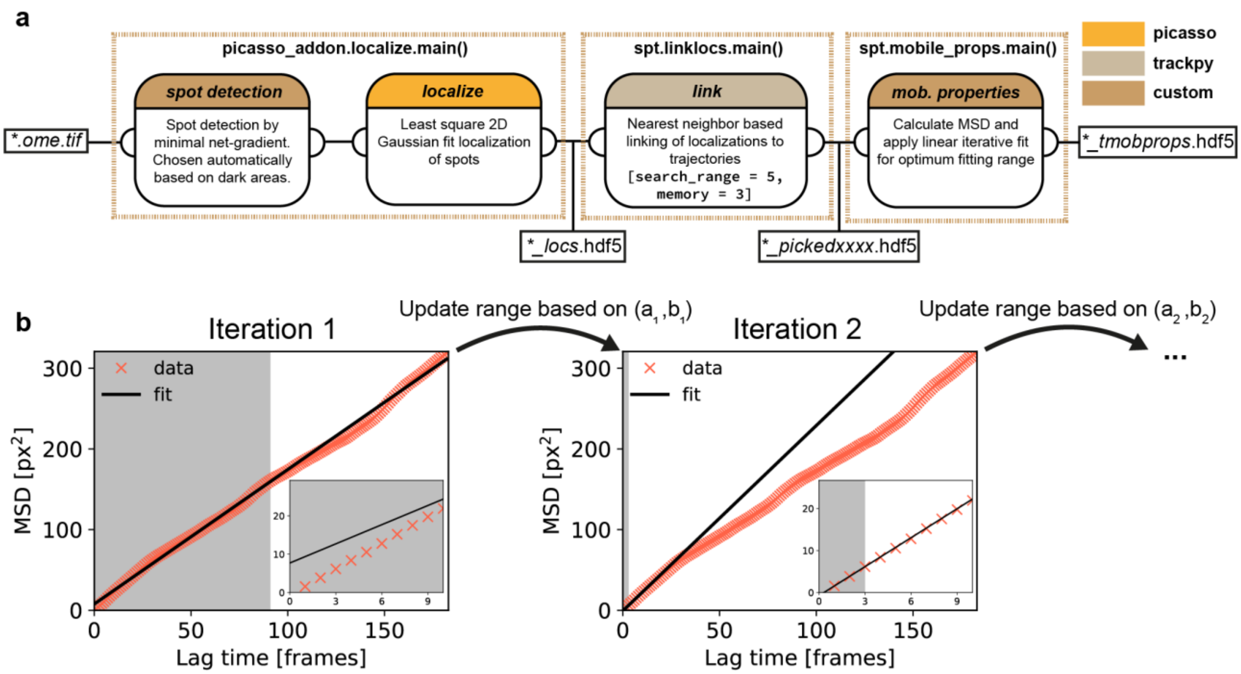


**Supplementary Figure 10. Data analysis workflow - mobile.** (**a**) Data analysis workflow from raw movies (*.ome.tif) to final result (*_tmobprops.hdf5). Dashed boxes indicate employed main functions - e.g. picasso_addon.localize.main() - of the *picasso_addon* and *spt* python package. Rounded boxes illustrate provided functionalities of each main function. The header color code indicates from which python package the functionality was adapted (e.g. *picasso, trackpy* or *custom* for extended functionalities within *picasso_addon* or *spt*). The text within the rounded boxes gives a short description of the provided functionality and parameters (brackets) for execution of the main function. For all evaluations of mobile origami a search_range value of 5 and a memory value of 3 was used (please refer to http://soft-matter.github.io/trackpy/v0.4.2/generated/trackpy.link.html#trackpy.link). The small boxes branching of the main flow represent which files are saved during execution. Please visit the links provided in the section "**Image processing & single particle tracking analysis**" for further information. (**b**) We followed a linear iterative fitting procedure of the individual MSD curves as proposed by Michalet et al.[7] to find the optimum fitting range. For the following description we define the  total trajectory length as $N$ and the maximum lag time $l$ up to which the MSD curve is fitted as $N_p$. In every step we fit the MSD with the linear fit model MSD$(l) = a \cdot l + b$  up to $N_p$. For the first iteration we set $N_{p,1} = 0.125 \cdot N$ (nearest integer) and perform an unweighted least square fit giving $(a_1, b_1)$. The left panel shows the fitting result of the first iteration. The grey area indicates the fitting range as given by $N_{p,1}$. The zoom-in illustrates poor fitting of the MSD values for short lag times $l$ which constitute the MSD values of lowest (statistical) uncertainty[8]. For the next iteration we hence update the fitting range as given by $N_{p,2}$ using the rule $N_{p,2} = 2 + 2.3(b_1/a_1)^{0.52}$ (rounded integer). The right panel shows the fitting result of the second iteration with an updated $N_{p,2}$ of 3. The zoom-in indicates that now the (low uncertainty) MSD values for short lag times $l$ are fitted well hence leading to a more precise determination of the diffusion constant. We usually observe a fast convergence in $N_p$ after 2 or 3 iterations and we only allowed up to five iterations.

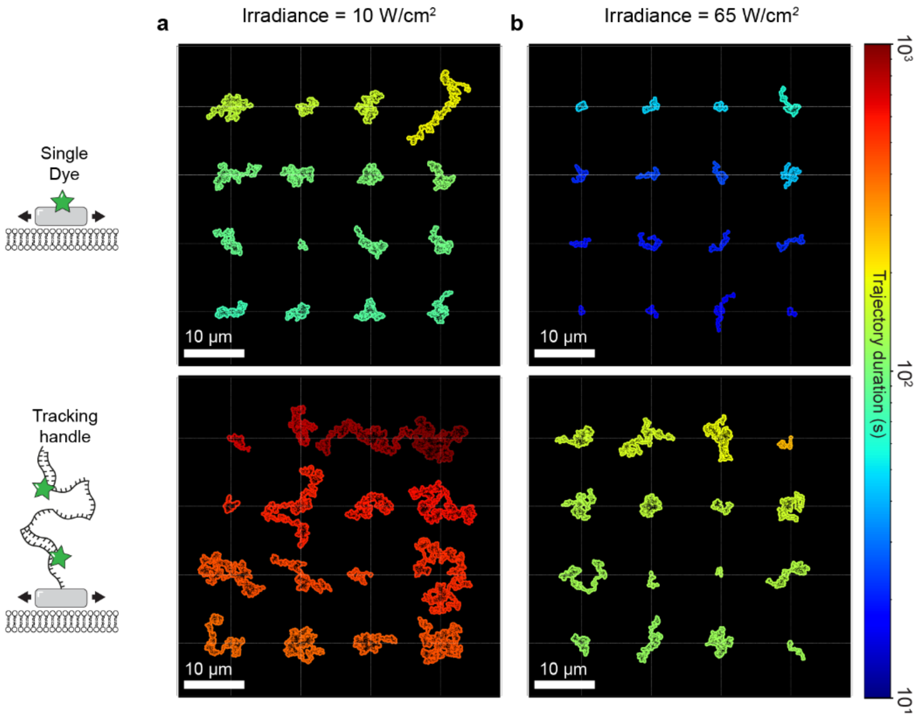


**Supplementary Figure 11. Longest trajectories at varying irradiances.** The 16 particle trajectories of longest durations of SD origami (top) and TH origami (bottom) floating on SLBs analogous to **Figure 4**a but measured with (**a**) lower irradiance and (**b**) higher irradiance. Longest observed TH trajectory duration (top right trajectory) for an irradiance of 10 W/cm$^2$ was ~ 30 min and ~ 4 min for 65 W/cm$^2$, respectively.

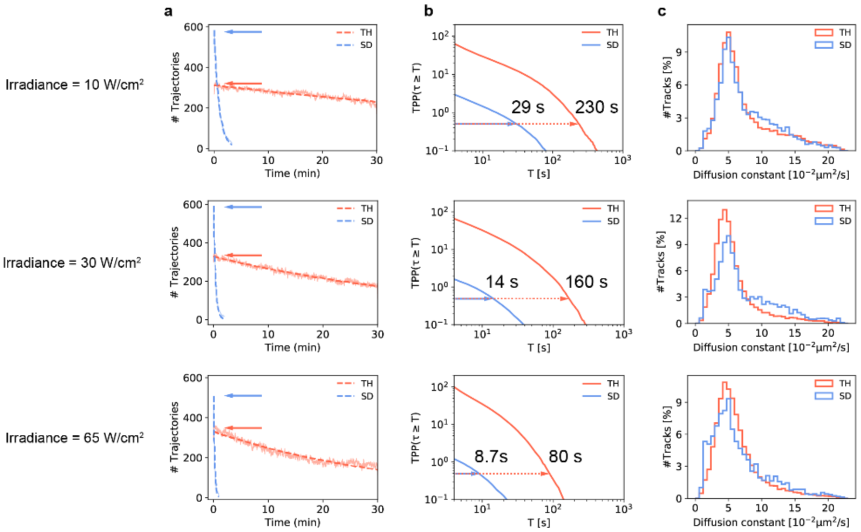


**Supplementary Figure 12. Tracks, TPP and diffusion constant at varying irradiances.** (**a**) Number of trajectories per frame analogous to **Figure 4**b. (**b**) Average number of tracks per origami $TPP(\tau_n \geq T)$ analogous to **Figure 4**c. (**c**) Diffusion constants as obtained by linear iterative fitting of the individual MSD curves analogous to **Figure 4**f. Floating TH origami results are indicated by orange color, floating SD origami results are indicated by blue color. Rows represent results as obtained for varying irradiances.

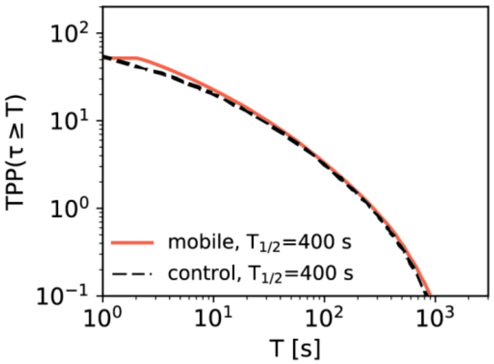


**Supplementary Figure 13. Control of mobile analysis pipeline on static DNA origami.** In order to verify our approach of TPP calculation for the mobile case (see section 'SPT of DNA origami on Supported Lipid Membranes' in main text) we reanalyzed an immobilized TH origami sample using the mobile analysis pipeline described in **Supplementary Figure 10**. The orange curve displays the corresponding TPP results. The black dashed line displays the TPP results as obtained via the immobile analysis pipeline described in **Supplementary Figure 2** and serves as a control**.** Both curves evolve almost identically and yield the same average TPP of ~50 and a $T_{1/2}$ value of 400 s. This confirms our approach for the mobile case to normalize the total number of detected trajectories to the number of trajectories (i.e. particles) in the first frame of the data set.

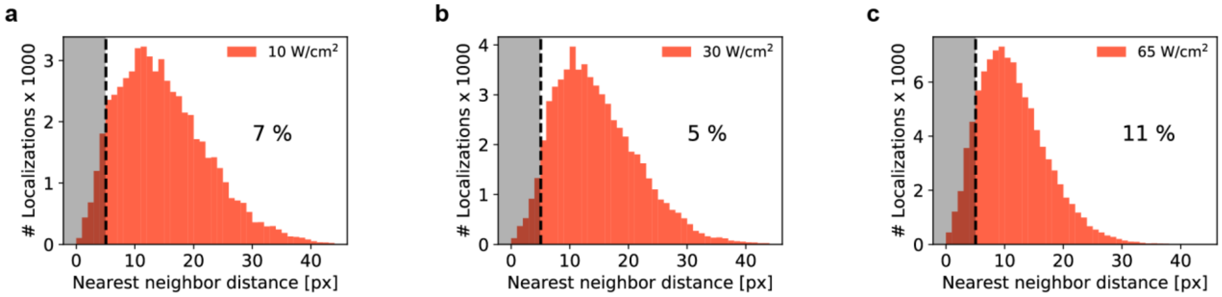


**Supplementary Figure 14. Linking and particle density**. (**a**) The data shown refers to TH origami measurements at an irradiance of 10 W/cm$^2$. It shows the nearest neighbor distance distribution between all localizations corresponding to the same frame of the recorded movie. The histogram represents the total distribution as calculated for each of the first 100 frames of the movie. The black dashed line indicates the range used as parameter for the nearest neighbor based linking algorithm (search_range in **Supplementary Figure 10**). 7 % of all nearest neighbor distances between localizations of one frame lie within the search range of the linking algorithm, potentially impairing its linking ability and thus leading to trajectories of shorter duration when compared to the immobilized samples. (**b,c**) Same as (a) but at an irradiance of 30 W/cm$^2$ and 65 W/cm$^2$, respectively.

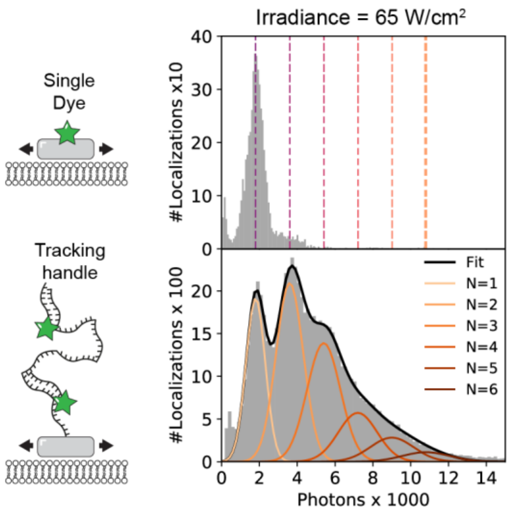


**Supplementary Figure 15. Photons counts - SD vs TH mobile.** Number of photons detected per localization for mobile SD origami showing a unimodal distribution (top). Number of photons detected per localization for mobile TH origami showing a multimodal distribution (bottom). A fit (black) consisting of the sum of 6 Gaussians (orange) was applied to analyze how many emitting imagers were present over time. The dashed lines in the top panel indicate the fit centers as obtained for the individual Gaussian functions of the TH origami (bottom). Both histograms refer to localizations from the central circular region of the FOV (diameter = 200 px) of the first 40 frames, 1000 frames of the measurement for SD origami and TH origami, respectively.

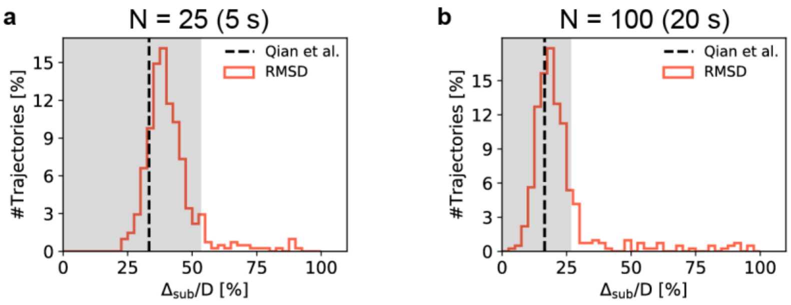


**Supplementary Figure 16. RMSD histograms for N=25 & N=100.** (**a**) RMSD distribution of $D_{sub}$ to $D$ of all trajectories exceeding 120 s split into subtrajectories of 5 s (red). RMSD was normalized to $D$ and should hence be close to the theoretical limit[8] (black dashed line) if the TH origami are subject to a time-invariant Brownian motion. Grey area indicates deviation of less than 60% to the theoretical limit. (**b**) Same as (a) but with a subtrajectory duration of 20 s.

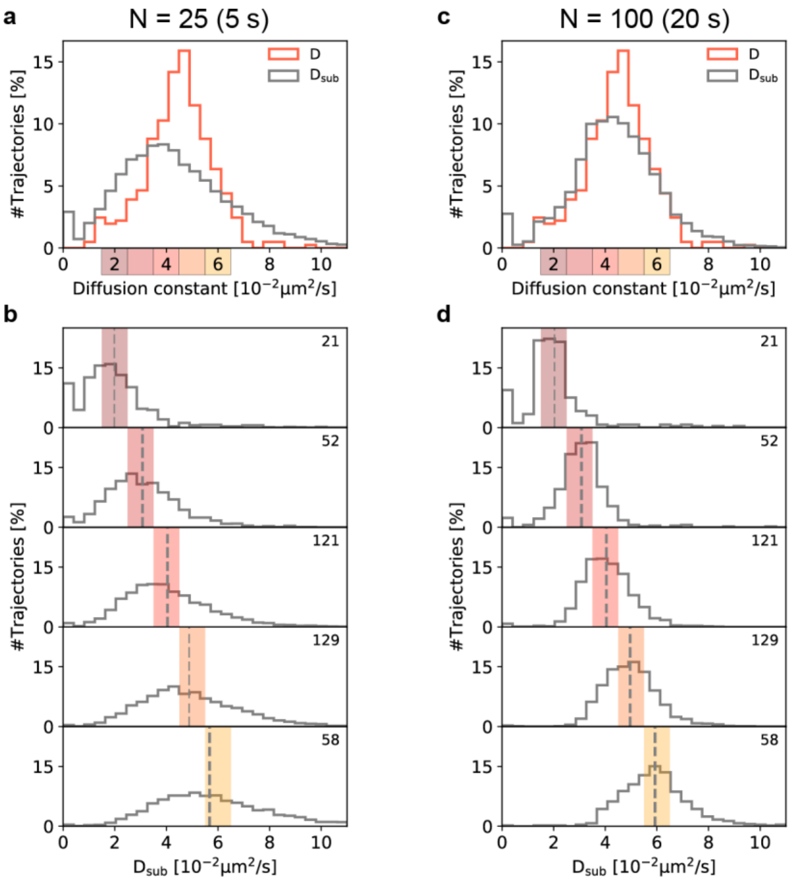


**Supplementary Figure 17. D vs. $D_{sub}$ for N=25 & N=100.** (**a**) Total distribution of $D$ and the corresponding subtrajectory diffusion constants $D_{sub}$ for subtrajectories of duration 5 s. (**b**) We selected five subsets of trajectories yielding a value of $D$ within the ranges 1.5 - 2.5, …, 5.5 – 6.5 µm$^2$/s (colored boxes in a) and plotted the corresponding $D_{sub}$ distribution. The mean value of $D_{sub}$ (grey dashed) agrees well with the selected central $D$. The top right number indicates how many trajectories were part of the selected subsets in $D$. We observed an average broadening factor of $D_{sub}$ to $D$ of 6x (theoretical: 6.7x, please refer to Eq. 1 in the main text). (**c**) Same as (a) but with a subtrajectory duration of 20 s. (**d**) Same as (b) but with a subtrajectory duration of 20 s. We obtained a broadening factor of 4x (theoretical: 3.4x).

### Supplementary Tables

#### Supplementary Table 1 | Imaging parameters

| Figure | Sample | Imager concentration (nM) | Imaging Buffer | Temperature (°C) | Irradiance (W/cm$^2$) | Frames (wrt. Irradiance) |
|---|---|---|---|---|---|---|
| 1d<br>2a,c,e,h,i | SD origami, static | - | L | 21 | 30 | 600 |
| 1e<br>2b,d,f,g,h,i<br>SI_Fig. 4 | TH origami, static | 40 | L | 21 | 30 | 9,000 |
| 3b, left panel | SD origami, static | - | L | 21 | 10<br>30<br>65 | 2,000 (3x FOVs)<br>600 (3x FOVs)<br>300 (3x FOVs) |
| 3b, left panel<br>3c, left panel<br>SI_Fig 7<br>SI_Fig 13 | TH origami, static | 40 | L | 21 | 10<br>30<br>65 | 9,000<br>9,000<br>9,000 |
| 3b, right panel | SD origami, static | - | POCT | 21 | 10<br>30<br>65 | 9,000<br>9,000<br>9,000 |
| 3b, right panel<br>3c, right panel | TH origami, static | 40 | POCT | 21 | 10<br>30<br>65 | 54,000<br>18,000<br>18,000 |
| 3d | TH origami, static | 5, 10, 20, 40<br>(4 samples) | L | 21 | 10<br>30<br>65 | 9,000<br>9,000<br>9,000 |
| 3e | TH origami, static | 5 nM | L | 21, 23 | 10<br>30<br>65 | 9,000<br>9,000<br>9,000 |
| 3f | TH origami, static | 40 nM | L, B<br>(2 samples) | 21 | 10<br>30<br>65 | 9,000<br>9,000<br>9,000 |

| Figure | Sample | Imager concentration (nM) | Imaging Buffer | Temperature (°C) | Irradiance (W/cm$^2$) | Frames (wrt Irradiance) |
|---|---|---|---|---|---|---|
| 4a,b,c,f,g<br>SI_Figs. 11, 12, 15 | SD origami, diffusing on SLB | - | L | 21 | 10<br>30<br>65 | 1,000 (3x FOVs)<br>600 (3x FOVs)<br>300 (3x FOVs) |
| 4 (all)<br>5 (all)<br>SI_Figs. 11, 12, 14 15, 16, 17 | TH origami, diffusing on SLB | 40 | L | 21 | 10<br>30<br>65 | 9,000<br>9,000<br>9,000 |
| SI_Fig 5a | SD origami, static | - | B (4x samples),<br>L (1x sample) | 21 | 10<br>30<br>65 | 2,000 (3x FOVs)<br>600 (3x FOVs)<br>300 (3x FOVs) |
| SI_Fig 5b | TH origami, static | 40 | L (2x samples) | 21 | 10<br>30<br>65 | 9,000<br>9,000<br>9,000 |
| SI_Fig 6 | TH origami, static | 40 | POCT | 21 | 10 | 54,000 |
| SI_Fig. 8 | 1DS origami | 5, 10, 20<br>(3 samples) | B | 21, 23 | 10 | 9,000<br>(6x) |
| SI_Fig. 9 | 1DS origami | 5, 10, 20 | B, L<br>(2x3=6 samples) | 21 | 10 | 9,000<br>(6x) |

**Supplementary Table 2 | Used DNA oligonucleotide sequences as labels**

| **Name (oligo length)** | **Docking strand sequence (5' – 3')** | **Imager sequence (5' – 3')** | **Experiment** |
|---|---|---|---|
| TH (54 bp) | TT | GAGGAGGA-Cy3B | All TH experiments |
| SD (5 bp) | TT TTT-Cy3B | - | All SD experiments |
| 1DS (8 nt) | TT TCCTCCTC | GAGGAGGA-Cy3B | lbFCS series in SI_Fig. 8 & 9 |
